# A generalized resource-allocation growth law reveals mechanisms of differential responses to intermittent androgen deprivation therapy in prostate cancer

**DOI:** 10.64898/2026.08.25.747066

**Authors:** Tin Phan, Heyrim Cho, Kyle Nguyen, Jeffrey West, Clay Prater, Puni Jeyasingh, Elizabeth Duke, Kevin Flores, Alan H. Bryce, Alan S. Perelson, Yang Kuang

## Abstract

Resource-limited growth frameworks often developed independently. Here, we derived a growth law by assuming a cell balancing internal competing resource demands and generalizing the growth signal through a shape parameter h, the trade-off exponent. In Liebig’s single-limiting-resource regime, it recovers Droop’s cell quota at h = 1 and approaches Prater’s growth efficiency as h grows large. h determines how sharply growth declines when allocation departs from optimal, thereby shaping populations evolution. Cyclic environmental stress tests such growth laws. Hence, we built a model from this growth law and applied it to prostate cancer growth in patients undergoing cyclic intermittent androgen deprivation therapy. We showed the model recapitulated longitudinal prostate specific antigen (PSA) and serum androgen measurements from 71 prostate cancer patients using a nonlinear mixed-effects framework (pooled and median individual R^2^ ≈ 0.9 for PSA and ≈ 0.8 for serum androgen). h is well-constrained by the data and separates treatment outcomes (success vs. failure, p = 0.003). The results predict that failing cases have cancer that can adapt to a wide range of growth conditions with minimal cost. A simplified model recapitulated PSA dynamics from 32 prostate cancer patients on cyclic adaptive therapy (R^2^ ≈ 0.8). Estimates of h for castration resistant cancer in adaptive cohort were similar to failure cases in the intermittent cohort. We found no statistical differences in estimated parameters between the adaptive vs. standard of care arms, suggesting that adaptive schedule may drive the differential outcome. The results support this growth law, providing a foundation for resource-limited growth across biological systems.

**Relevance:** Droop’s cell-quota model, Scott-Hwa proteome allocation, Sterner-Elser stoichiometry and Prater’s growth efficiency, and Liebig’s law of the minimum all seem like different frameworks for resource-limited growth. We derive a single growth law from the idea that cells balance competing resource demands, with growth slowing when this balance is off, and we show that it connects these frameworks. Applied to trials of intermittent and adaptive androgen deprivation therapy, the law predicts that failing tumors pay little growth cost when they adapt, so resistance arises easily. It further suggests adaptive treatment schedules drive outcome instead of cohort differences. The same law connects ideas from ecology, microbiology, evolution, and medicine, offering a quantitative framework for how living systems grow under limited resources.

## Introduction

Resource-limited growth spans across biology. Every organism draws resources from an environment often beyond its control and allocates them among competing functions. The earliest work focused on the environment. Sprengel and Liebig showed that yield is determined by the nutrient in shortest supply relative to demand, a principle known as the law of the minimum (1). Blackman later expressed an analogous idea for physiological rates (2) and Monod then drew on the saturating form of enzyme kinetics to show that bacterial growth depends similarly on the concentration of the limiting substrate (3). By the middle of the century, a coherent picture began to emerge, growth increases with the availability of external resources but eventually saturates, and the resource in shortest supply sets the rate. For many organisms, however, this picture is incomplete. Algal growth rate is set by an internal nutrient quota, so cells can continue dividing after the external nutrient has been exhausted (4, 5). Droop resolved this discrepancy by considering resource availability inside the cell. In his model, growth depends on the cell quota (Q), the amount of the limiting resource stored within the cell, relative to a subsistence quota (q), below which growth ceases (6, 7). Uptake and growth are separate processes, coupled through an internal storage. Tilman showed that the same internal-storage logic helps determine which species persist when several compete for scarce nutrients (8).

The field of ecological stoichiometry later emerged, building on Redfield’s observation that marine plankton have an average atomic N:P ratio of approximately 16:1 (9). It treats elemental composition as both a constraint on organismal function and a signature of that function. The growth-rate hypothesis turned this connection into a testable prediction: rapid growth requires greater allocation to phosphorus-rich ribosomal RNA (10, 11). More recently, Prater et al. recast stoichiometric limitation as an imbalance between resource-use efficiencies (12). Bacterial physiology also shifted the focus from nutrient availability to the allocation of cellular machinery. Building on the observation that bacterial composition tracks growth rate more closely than it does the identity of the growth medium (13), Scott, Hwa, and colleagues showed that growth rate is determined by two opposing linear constraints on the tri-allocation of the proteome (14, 15).

These frameworks use different independent variables, units, and disciplinary vocabularies, and they arose mostly in separate literatures. Yet they share the same underlying intuition: an organism must allocate a finite internal supply among competing functions, and its performance declines when that allocation fails to meet their demands. This is a trade-off of the kind that life-history theory has long regarded as fundamental to organismal performance (16, 17). Others have shown that responses to multiple resources span a continuum from fully substitutable to strictly complementary (18–21). This common intuition suggests a structural connection among all of these growth laws.

Here we derive a single growth law that connects Droop’s cell-quota model, Scott-Hwa proteome allocation, Sterner-Elser stoichiometry and Prater’s growth efficiency within one construction. The derivation starts from the resource allocation framework of Scott-Hwa (14), extends its interpretation to Droop’s cell quota, and then generalizes by introducing a trade-off exponent parameter, h, which measures how sharply growth declines when allocation departs from optimal. The law reduces to Droop at h = 1, asymptotically converging into Prater’s growth form for large h, all within Liebig’s single limiting resource regime. We then apply the framework to intermittent and adaptive androgen deprivation therapy for prostate cancer (22, 23), settings with rich longitudinal data and a well-defined clinical endpoint and show that the trade-off exponent is the dominant outcome-associated parameter in the fitted model.

## Results

### A generalized resource-allocation growth law

Consider a cell that splits its internal resource between machinery serving growth and survival (Fig. 1A). We use “machinery” as shorthand for a proteomic sector in the sense of Scott et al. (14), although the derivation does not depend on the actual molecular make-up of the sectors. Let r denote the allocation ratio between these two functions and λ the specific growth rate. Following Scott et al. (14), the growth rises linearly with allocation r, λ_g_ = c_1_(r − r_0_), in resource-abundant environment, while in the stressed or resource-scarce regime, survival capacity is necessary and λ_s_ = c_2_(r_m_ − r). Here r_m_ and r_0_ denote the maximum and minimum possible value for r, and c_1_and c_2_ the respective slopes of the two branches. We refer to the rising line as the growth-limited branch (λ_g_) and to the falling line as the survival-limited branch (λ_s_). A single internal budget must serve both demands at once, so the realized growth rate is set by whichever sector is scarcer. This is Liebig’s law of the minimum applied to internal allocation, which makes the realized growth rate follows the lower of the two branches. Hence a cell that regulates allocation to maximize growth will operate at the intersection of the two branches (Fig. 1B) (24). Thus, setting the two branches equal (λ_g_ = λ_s_), solving for r, and substituting back gives the realized growth rate λ,

**Fig. 1.**
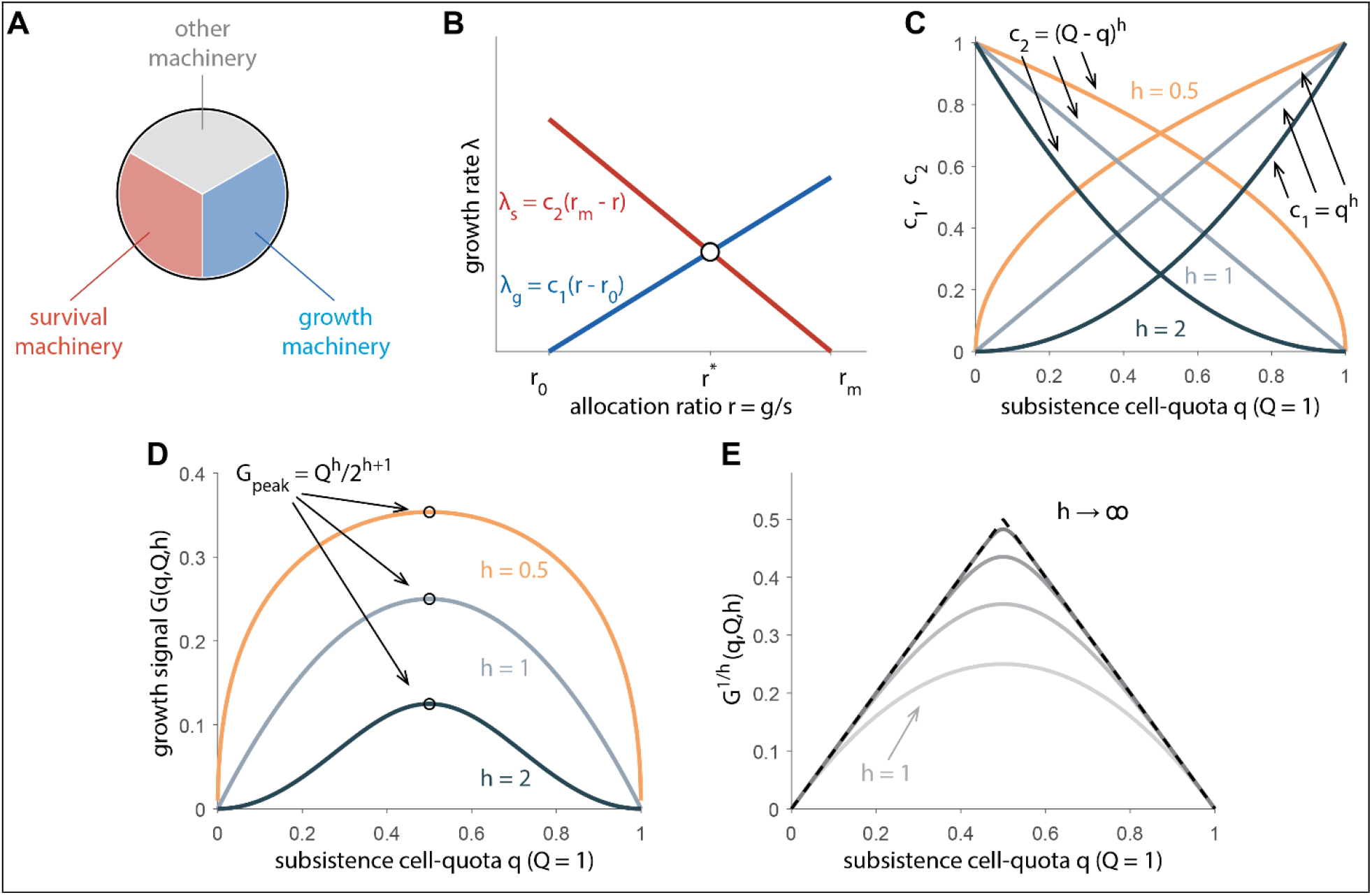
Derivation and implication of the generalized growth law. (**A**) A cell splits internal resource among growth machinery (blue), survival machinery (red), and machinery for other functions (grey). (**B**) In resource-abundant conditions, growth rate rises linearly to the growth-to-survival allocation ratio r, i.e., λ_g_ = c_1_(r − r_0_). In resource-scarce conditions, growth rate decreases linearly to r, i.e., λ_s_ = c_2_(r_m_ − r). The intersection gives the realized growth rate λ = (r_m_ − r_0_)c_1_c_2_/(c_1_ + c_2_). (**C**) Dependence of the growth and survival coefficients on the subsistence cell quota. The growth coefficient c_1_increases with subsistence cell-quota q, c_1_= *q*^*h*^, whereas the survival coefficient c_2_ decreases with q. The trade-off exponent h determines the curvature of these relationships, shown for h = 0.5, 1, 2. (**D**) The shape of the growth signal G(q, Q, h) for h = 0.5, 1, 2. (**E**) The generalized growth converges to Prater’s growth efficiency (dashed line) for large h.

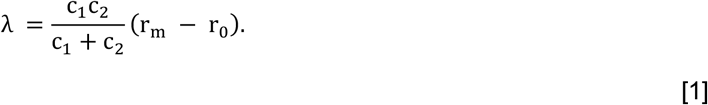

The factor 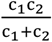 in Eq. 1 is proportional to the harmonic mean of the two slopes, small whenever either slope is small. This implies growth is slowed if either growth or survival are not optimized. Next, motivated by Droop’s cell-quota relationship 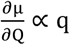, we assume that the marginal benefit,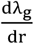, of allocating more resource toward growth machinery, in a resource-abundant environment, scales with the minimum amount of that resource the cell requires in order to grow q (25). Hence, c_1_∝ q, where q is the subsistence cell quota – the minimum amount of the limiting internal resource required for an organism to grow. This implicitly assumes that q is a function of the machinery allocated for growth: a high-q organism is one that has invested heavily in growth machinery, allowing it to grow rapidly in a resource-rich environment, and vice versa. However, more growth machinery is also more expensive to maintain, especially in resource-limited environment. Consequently, we assume that the marginal cost of losing survival capacity in a resource-scarce environment scales with the available surplus above subsistence cell quota, e.g. c_2_ ∝ Q − q, where Q is the cell quota, the current internal pool of the limiting resource contained per cell, for example, cellular phosphorus in a phosphorus-limited alga. The proportionality constants are the same for both relationships, since they represent the same conversion of machinery to growth. We define μ_0_ as the baseline growth rate which absorbs the proportionality constant r_m_ − r_0_. This gives a trade-off version of Droop’s growth (25):

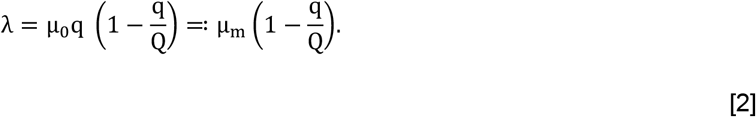

Eq. 2 contains a novel structural feature that is not part of the classical Droop’s form. The maximum growth rate is not an independent parameter but is set by the machinery allocated for growth, i.e., μ_m_ = μ_0_q. However, we can take this further and relax the linearity assumption since this derivation holds for any monotone function of c_1_ and c_2_. We introduce a trade-off exponent h, giving c_1_∝ q^h^ and c_2_ ∝ (Q − q)^h^ (Fig. 1C). The generalized growth rate takes the form:

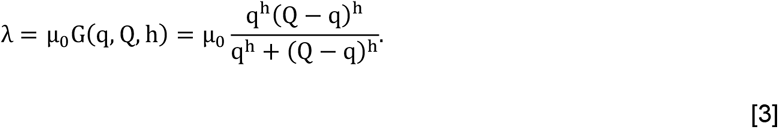

The generalized growth form G(q, Q, h) in Eq. 3 is unimodal in the subsistence quota and peaks when q = q^optimal^ = Q/2. It also recovers the trade-off Droop’s form (Eq. 2) when h equals 1, i.e., the denominator of Eq. 3 collapses to q + (Q − q) = Q, leaving G = q(Q − q)/Q = q(1− q/Q). Throughout, h > 0 and G is evaluated on the biologically feasible domain 0 < q ≤ Q, so that Q − q represents available surplus resource, which may be required for other cellular processes than growth or survival. Biologically, h describes how sharply growth declines when q drifts off the balanced midpoint (Fig. 1D). Large h (convex curve; h > 1, Fig. 1D) implies a heavy cost for over-investing in either growth or survival, with diminishing marginal returns to growth or survival. Conversely, small h (concave curve; h < 1, Fig. 1D) implies tolerance to a wide range of allocation choices. The exponent therefore sets marginal returns, e.g., doubling the growth allocation more than doubles its contribution when the exponent exceeds 1, and less than doubles it when the exponent is below 1. Increasing marginal cost is expected when growth and survival machinery compete for the same resource. On the other hand, decreasing marginal cost is expected when these functions have partly decoupled, so that survival capacity is acquired at little cost in growth, e.g. an alternative signaling route in tumors. This implication runs parallel to theories on evolutionary trade-off such as Pareto optimality (26) or Fisher’s geometric model (27). We also see that the peak value 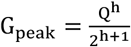 links the maximum growth to the cell quota Q based on the trade-off exponent h, e.g., μ_m_ is a linear function of Q with slope h on log scale. For very large h, μ_0_G^1/h^ asymptotically converge to Prater et al. (12), which posits that an organism’s growth rate declines linearly from its maximum as the total imbalance in resource use increases under nonoptimal conditions:

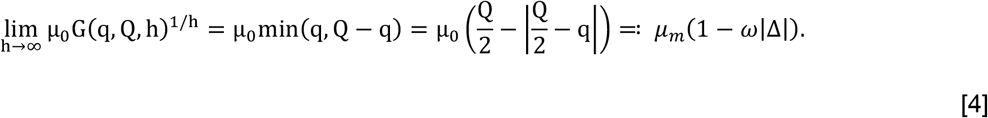

In Eq. 4, 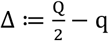 is the difference between the current allocation q and the optimal allocation q^optimal^ = Q/2, with μ_m_ ≔ μ_0_q^optimal^ = μ_0_(Q/2) the maximum growth rate at optimal allocation, and ω ≔ 2/Q the scaling constant. The exponent h maps the growth G^1/h^ between Droop’s (h = 1) to the Prater’s growth efficiency as h → ∞ (Fig. 1E).

When looking at population growth, the subsistence quota q can be considered a trait, e.g., allocation strategy, that evolves under selection using the breeder’s equation (28) (see S3 Text),

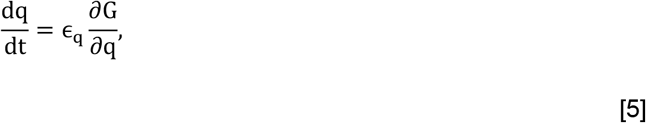

where ϵ_q_ is an effective evolutionary rate constant and 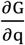 is the selection gradient (29, 30). The gradient is positive when q < Q/2 and negative when q > Q/2, that is selection always pushes q toward the optimal midpoint. The closed form for 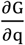 follows from Eq. 3 (see S2 Text).

### Application to intermittent androgen deprivation therapy in prostate cancer

Prostate cancer cells depend on androgen signaling for growth and survival (31). When androgen is suppressed, this inhibits the growth of the androgen-sensitive prostate cancer cells, which usually form the majority of the cancer cell population prior to treatment (32, 33), and can lead to remission (34). However, continuous androgen suppression ultimately selects for cancer cells that require less androgen, or that can proliferate despite castrate androgen levels, while also imposing substantial negative side effects on patients (35). This has motivated research into intermittent androgen deprivation therapy (IADT), in which patients alternate on and off treatment with the goal of improving quality of life and potentially delaying treatment resistance (36). Here, androgen can be considered as the limiting resource for prostate cancer cells, one that they must allocate their molecular machinery to use efficiently (37). This cyclic treatment over several years provides an example of our framework, since each cycle is a controlled perturbation of the limiting resource.

We developed a model for cancer growth based on the generalized growth Eq. 3 and previous work (38– 40) (Fig. 2). The model tracks cancer cell population x, subsistence androgen quota q – the intracellular androgen concentration cancer cell requires for grow, intracellular androgen concentration Q, serum androgen concentration A, and serum prostate specific antigen (PSA) concentration P. The cancer mass equation is 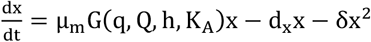, where d_x_ and δ are the per capita and density-dependent death rates and G(q, Q, h, K_A_ ) is the normalized growth form in Eq. 3, e.g.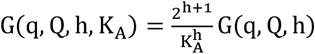. Serum androgen A is produced by the testes and adrenal gland at an effective rate γ_e_ with a production capacity K_A_, and decays at per capita rate d_A_. Applying the law of mass conservation, intracellular androgen Q equilibrates with serum androgen at rate v and dilutes as the cell grows at rate μ_0_G(q, Q, h, K_A_). The androgen subsistence quota q evolves toward q^optimal^ according to breeder’s equation (Eq. 5). Here, the function G represents proliferation rather than net growth (38, 40). Accordingly, androgen deprivation acts only on the proliferation term, so when Q falls below q, proliferation ceases (G is evaluated for 0 < q ≤ Q, see S3 Text), whereas cancer death terms d_x_ and δ are independent of q and Q. Thus, androgen deprivation stops proliferation and cancer cell turnover leads tumor regression. Serum PSA is produced by benign tissue, or non-cancerous prostate cells, at rate bA and cancer cells at per capita rate σ_c_H(G). The drug effect is represented by D with a wash-out time τ_w_, while the progressive loss of androgen recovery (i.e., a decrease of K_A_) over each treatment cycle (41, 42) is described by the homeostatic androgen decay rate κ. Here the production capacity is the patient eugonadal androgen target, obtained from the observed androgen record (S3 Text). This applies the growth law to a dynamic-environment setting. Finally, u(t) is the treatment indicator (u = 1means treatment is on; u = 0 means treatment is off). The model equations are Eqs. [6] – [15] with its full derivation and the definitions and units presented in S3 Text and Table S1. A reduced model with simplified androgen dynamics is derived and used to fit the adaptive therapy cohort from the Moffitt Cancer Center (23) (see S4 Text). The parameter ranges are the same for the two models.

**Fig. 2.**
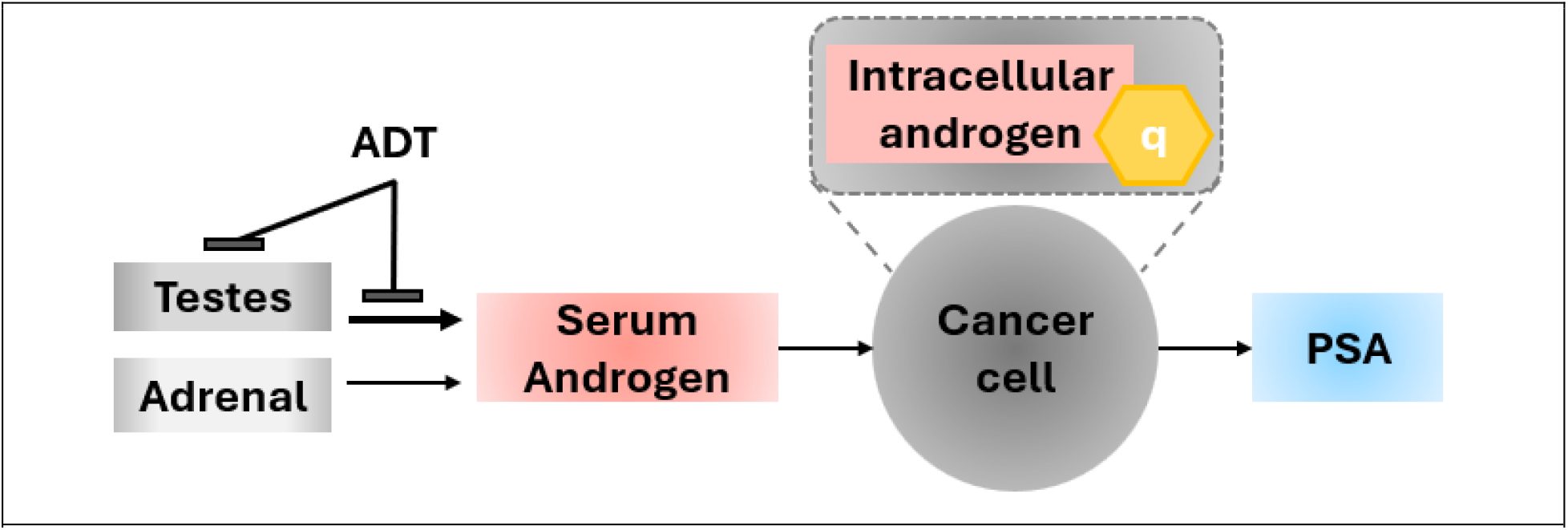
A simplified schematics for the prostate cancer model. Serum androgen is produced by testes and the adrenal gland. Cancer cells uptake androgen to grow and produce PSA in the process. Androgen deprivation therapy suppresses the androgen production capacity of the testes and thus hinders cancer cell growth. Full model equations and their derivation are presented in S3 Text.

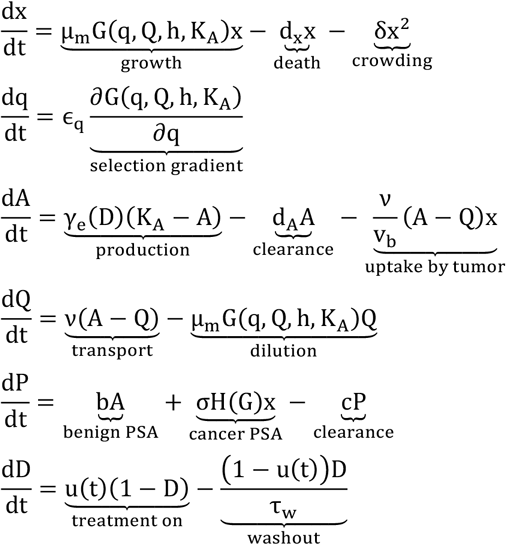

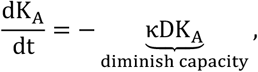

where

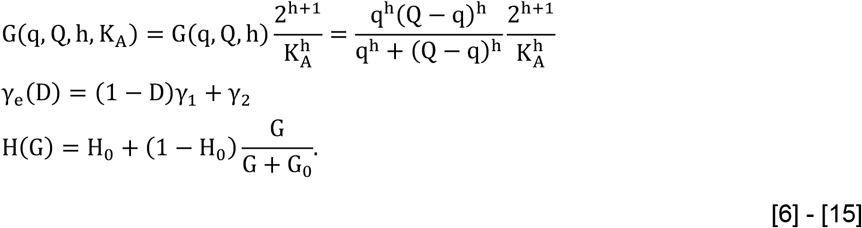

We use a nonlinear mixed effect modeling framework (Monolix, Lixoft 2024) to fit the model to longitudinal PSA and androgen data from 71 patients (22), totaling 6,114 longitudinal observations to estimate ten parameters: μ_m_ (max growth rate), h (trade-off exponent), ϵ_q_ (quota evolution rate constant), γ_1_(testicular androgen production rate), d_A_ (androgen clearance rate), f_C_ (Cancer-derived fraction of baseline PSA), d_x_ (per capita cell death rate), δ (density-dependent cell death rate), q_F_ (initial subsistence quota fraction), and κ (homeostatic androgen decay rate). Of these 71 patients, 56 completed the trial without biochemical failure and 15 experienced biochemical failure, defined by the original trial protocol as a sustained rise in serum PSA while on treatment, indicating loss of androgen sensitivity. We refer to these two groups as success and failure, respectively. The model recapitulates the major cyclic PSA and androgen dynamics across patients (selected results in Fig. 3A; full results in Fig. S1 and S2). The reduced model also recapitulates longitudinal PSA data from the adaptive cohort (Fig. S3). The *R*^2^ for model fit vs. observation for PSA and androgen are ∼0.9 and ∼0.8 for the Bruchovsky’s cohort, and ∼0.8 for the adaptive cohort (see Fig. S4 and S5).

**Fig. 3.**
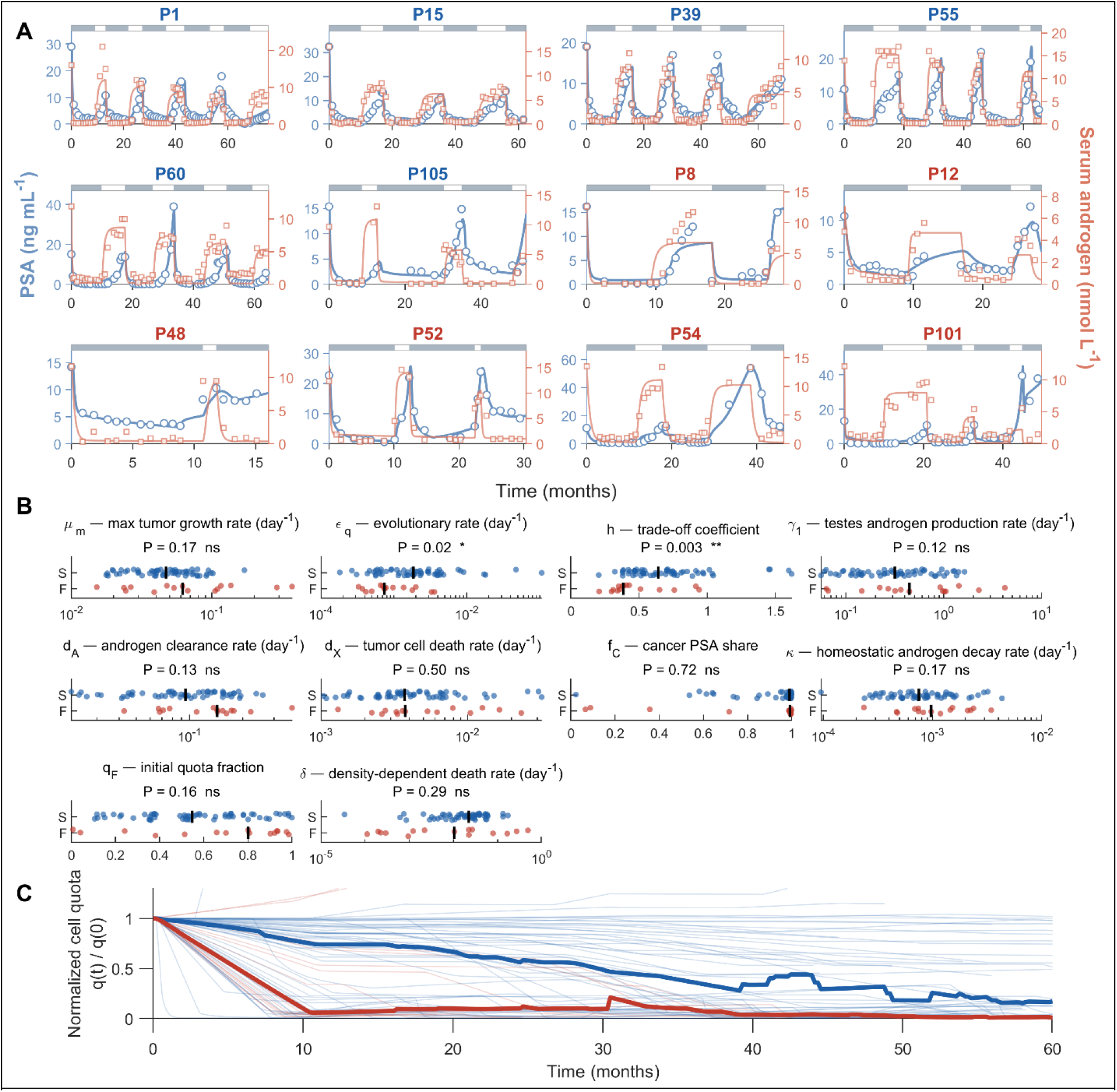
Model fit to Bruchovsky et al. data and analysis. (**A**) Model fit to longitudinal PSA and androgen data for 12 representative patients (full results in Fig. S1 and S2). Blue points/curves are PSA observations/model fit; Red points/curves are serum androgen observations/model fits. Grey bar on top indicates on treatment. White bar is off treatment. Blue subtitle indicates success, while red subtitle indicates failure. (**B**) Stratification of individual patients’ parameters based on the reported outcome (S: success; F: failure). Mann-Whitney U-test were used for statistical comparisons. With ten tests, the Bonferroni-corrected threshold for significance at α = 0.05 is 0.005. (**C**) Temporal variation of the normalized subsistence cell quota. Blue is the median of the successful cases, while red is the median of the failure cases.

### Treatment failure is associated with a lower growth trade-off

The dominant outcome-associated quantity and the only significantly different parameter after Bonferroni correction for ten parameter-wise tests (threshold: p = 0.005) is the trade-off exponent h (Fig. 3B), which is estimated with low population-level uncertainty at h = 0.61(95% CI 0.54 − 0.69, relative standard error (RSE) 6.4%), and the confidence interval excludes 1. The reference scale is set by the classical framework of 1 (Droop model) and h mostly determines the convexity of the trade-off, so values below 1 describe growth that is less sensitive to allocation imbalance than the Droop model. An important finding is that patients who fail therapy have lower h than patients who succeed (median 0.38 vs 0.64; p = 0.003). In addition, the reduced model fitted to 32 patients from the adaptive therapy trial at Moffitt Cancer Center also estimated a median trade-off exponent h of 0.44, statistically no different from the Bruchovsky failure group (p = 0.26). Since the adaptive trial only enrolled patients with castration resistant cancer, the same phenotype we expect to see in failure cases of the Bruchovsky trial, this supports the framework’s ability to estimate these parameters robustly across cohorts.

In this context, the competing demands captured by h are related to the machinery allocated to growth and survival under androgen deprivation. The seven patients whose estimates of h exceed 1 are all successes. The two regimes of trade-off exponent have opposite therapeutic consequences. Looking at the concavity of growth curve through the lens of antifragility theory (43), low values for h lead to a strongly concave growth curve (Fig. 1D) with a plateau-like peak whereby a cell population can experience larger variation in q without a severe penalty to growth. A large h means a sharper trade-off, where the growth falls steeply as allocation departs from balance, so the tumor cannot become resistant without losing growth, and the alternating treatment regime keeps forcing it to pay that cost. This is most clearly observed during treatment, where androgen deprivation reduces Q − q, which in turn reduces the growth rate. But the concave nature of the growth versus q curve means that the relationship between treatment and growth is convex, implying that higher dose values will be increasingly effective, especially when compared to small to moderate doses (e.g. doubling the dose gives more than double the effect (43)). For example, at the failure median h = 0.38, removing half of the available androgen leaves 77% of the growth signal. A small exponent therefore describes a tumor whose growth signal and the immediate PSA response are insensitive to partial suppression.

The smaller evolution rate constant ϵ_q_ in failure cases (Fig. 3B, p = 0.02 – not significant under Bonferroni correction) shows an unexpected trend: success patients have a faster cancer evolutionary rate compared to those failing therapy. However, the trajectory of the normalized androgen quota q(t)/q_0_ tells a different story (Fig. 3C). The net rate at which q changes is ϵ_q_ multiplied by 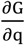, which is steep at low q when h < 1because it changes as q^h−1^(see Eq. S2). This means a tumor with small ϵ_q_ can adapt rapidly if h is small and q is far away from q^optimal^. Thus, despite the smaller evolutionary rate constant ϵ_q_, the normalized androgen quota trajectories in failing patients adapt rapidly during treatment transitions. This suggests failing tumors are not slow-evolving, they are fast-adapting yet weakly penalized when off-optimum.

### Eco-evolutionary tumor parameters stratify patients by treatment-failure risk

A Cox proportional-hazards model on the estimated parameters as predictors separates patients predicted to succeed from those predicted to fail treatment (Fig. 4). When evaluated in the full cohort of 71 patients (in-sample, Fig. 4A), 14 of the 15 failures fell in the high-risk group, giving a hazard ratio (HR) = 19.5 (95% CI 2.6–149), p < 0.001, and concordance index C = 0.88. Here HR is how much faster the high-risk group fails compared to the low-risk group at any given moment, and C is the chance the model correctly picks which of two random patients will fail first (C = 0.5 means random, C = 1.0 means perfect). Under leave-one-out (LOO) validation (Fig. 4B), the hazard ratio dropped to 9.0 (95% CI 2.0–39.9), p < 0.001, with C = 0.79. The decrease in model performance from in-sample to LOO is expected since with only 15 failures and 10 predictors, the in-sample model performance is inflated. The two curves separate clearly by month 20. The low-risk group stayed almost entirely failure-free across the 7-year follow-up, with only 1 of 36 patients reaching the failure endpoint (the rest were failure-free at the last follow-up). The widening confidence bands after roughly 50 months reflect the small number of patients remaining under follow-up that late.

**Fig. 4.**
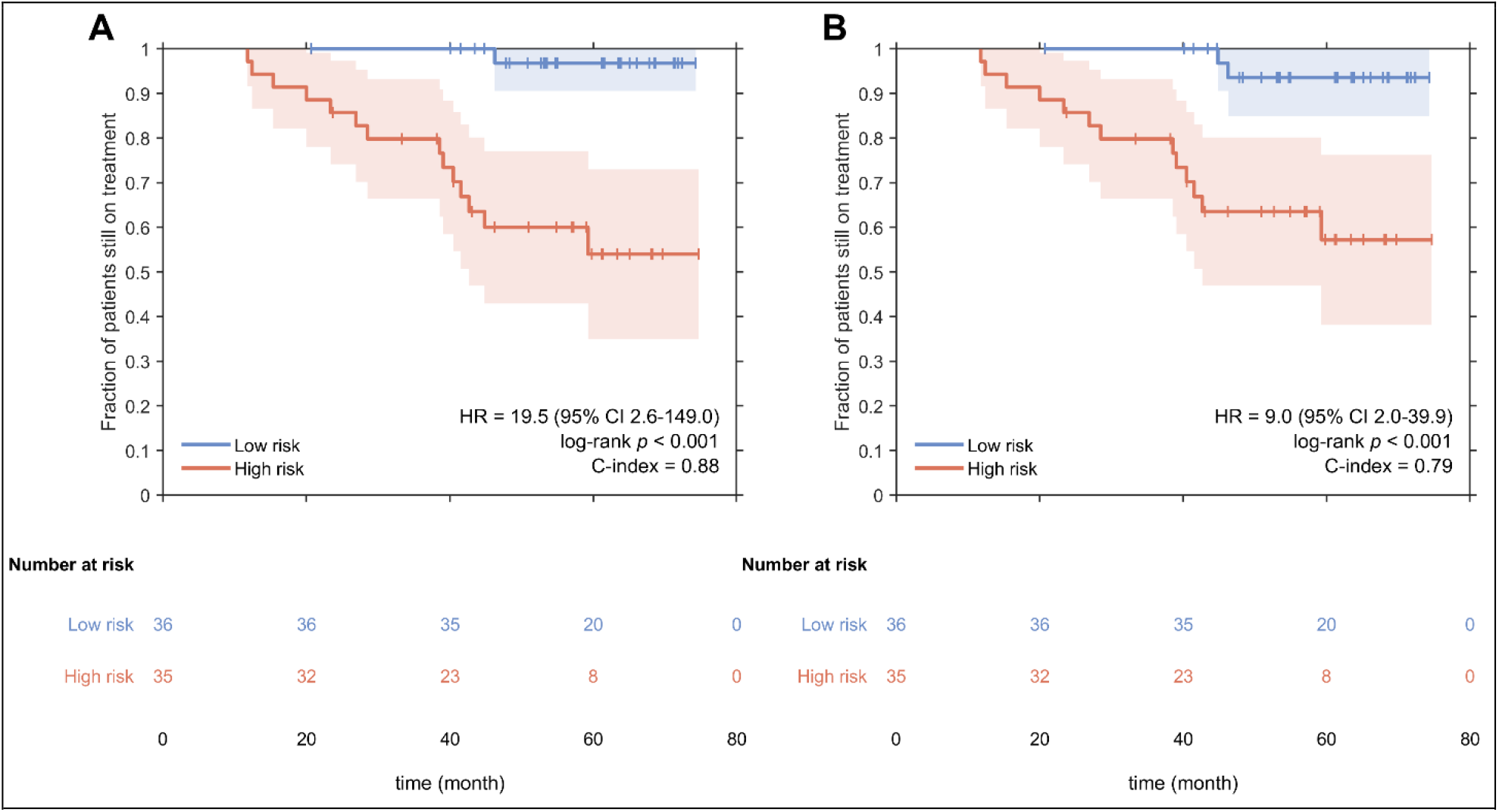
Patients stratified into low- and high-risk groups by a Cox model fit to parameters of individual patients. **(A) In-sample:** a Cox model was fit on all 71 patients using all parameters as predictors, and patients were split at the median predicted risk into low-risk (blue, n = 36) and high-risk (coral, n = 35) groups. **(B) Leave-one-out validation:** the same procedure was repeated 71 times, each time leaving out one patient and predicting that patient’s risk from the model fit on the other 70. Curves show the fraction of patients still on treatment over time, with shaded 95 % confidence bands. Small vertical ticks mark censoring, indicating patients whose last follow-up ended without treatment failure. Text in each panel reports the hazard ratio (HR) comparing the two groups with 95 % confidence interval, the log-rank p-value for the difference between the curves, and the concordance index (C-index), which ranges from 0.5 for random ordering to 1.0 for perfect ordering. Numbers below each panel show how many patients were still being followed at each 20-month checkpoint.

## Discussion

Biological life is shaped by what is feasible, but at what cost. This trade-off has been formalized many times in the resource-limited growth literature such as Droop’s cell quota (6), Liebig’s law of the minimum (1), Prater’s growth efficiency (12), and Scott-Hwa proteome allocation (14). Here we show that within Liebig’s single-resource regime, these frameworks share a hidden underlying structure governed by a single trade-off exponent h. This formalizes the biological trade-off in terms of increasing or decreasing the marginal benefit to allocating resources to survival relative to growth. This phenomenon is driven by the concavity, or convexity, principle that determines the effect of varying resource allocation, e.g. in response to treatment, on growth dynamics.

The growth form G(q, Q, h) generalizes continuously from Droop’s to Prater’s formulations and provides a unifying interpretation of these frameworks. Small h means a wide range of resource allocation strategies achieve nearly the same growth rate, while large h confines competitive strategies to a narrow window near the optimum (Fig. 1D). At h = 1(Droop), the selection gradient ∂G/ ∂q at fixed Q is (Q − 2q)/Q, which acts like a linear restoring force, analogous to an ideal spring (Hooke’s law). And similar to spring potential energy, the cost in growth is quadratic with deviation from the optimum (i.e., 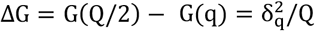 with δ_q_ = Q/2 − q). Thus, small imbalances are nearly free, while larger ones become disproportionately costly. As h grows larger, growth cost rises more steeply with deviation from the optimal allocation (see S2 Text). Prater’s growth efficiency (h → ∞) hence reflects a more tightly engineered organism whose growth and survival machinery are stoichiometrically locked. This implies application of Prater’s framework to multi-dimensional nutrient profiles would predict narrow, well-separated allocation niches – clusters of differentiated local strategies influenced by few nutrients that permit growth under a limiting environment, as the name “growth efficiency” would suggest.

On the other extreme, small h means little (h < 1) to no trade-off (h = 0) relative to the limiting nutrient. Biologically, this is the regime of an organism whose growth has been decoupled from the limiting resource. For prostate cancer, small h implies a phenotype that can either amplify the residual androgen signal during ADT via androgen receptor amplification or use an alternative signaling pathway. When the environment is perturbed so that an important resource is limited, selection can drive the organism toward the low h regime over time, with h itself potentially decreasing as the population adapts. Our clinical data analysis supports this picture. In the Bruchovsky cohort (hormone-sensitive disease under IADT), patients who fail therapy have a substantially lower trade-off exponent than those who succeed: median h is 0.38 in failures (n = 15) versus 0.64 in successes (n = 56), p = 0.003. We also fit the model to a cohort of patients in an adaptive therapy trial from the Moffitt Cancer Center (23) (reduced model and fits in Fig. S3A and S4 Text). The adaptive therapy cohort, comprised entirely of metastatic castration-resistant patients who had already progressed past first-line ADT and were treated with abiraterone (an inhibitor of androgen biosynthesis), has a similar median h of 0.44 (n = 32, IQR [0.39, 0.52]), statistically no different than the Bruchovsky’s failure cohort (p = 0.26). Across two independent cohorts and disease states, low h identifies the same biological phenotype: a cancer population that has weak dependence on androgen. The similarity in estimated h (medians 0.43 vs. 0.47, p = 0.88, Fig S3B) and all estimated parameters (Fig S3B) for the adaptive and standard-of-care arms in the adaptive cohort also support that the success of adaptive therapy is due to the treatment schedule and not intrinsic cancer difference.

By the same argument, a low trade-off exponent makes the dose-response relationship convex (Fig 1D), which implies partial doses buy disproportionately little suppression. This is the mirror image of the situation Strobl et al. analyzed for PARP-inhibitor maintenance in ovarian cancer, where a saturating dose-response made continuous dose modulation the better choice (44). We note this parallel without claiming that the same scheduling result applies here. Both trials used a single dose level that was either on or off, so we deliberately has no explicit dosing in our model. Moreover, the time to resistance depends on the gradual decline in q and on cumulative androgen suppression, neither of which can be inferred from the curvature of the instantaneous response alone. Resolving this question will require consideration of explicit drug dosing (45) and clinical trial that varies dose intensity. Nevertheless, the small trade-off value found here support previous work suggesting that adaptive therapy could work even if gaining resistance incurs a low cost for cancer (46).

The existing modeling literature on prostate cancer spans spatial and compartmental models of tumor growth and PSA kinetics (47–49), PSA-based prediction of intermittent-therapy responses (50), parameter identifiability and measurement selection (51–54), and treatment scheduling, including adaptive regimens with toxicity constraints (55, 56). Our framework complements these approaches by focusing on the form of the growth law itself. In principle, the trade-off exponent could be estimated by implementing our growth formulation within any of those frameworks. Furthermore, because the framework incorporates both androgen and PSA in a mechanistic way, it can be readily applied to trials such as A-DREAM (Alliance A032101) (57), where the endpoint combines PSA and androgen criteria. Although different trial designs may alter the limiting resource on different schedules, the exponent has the same interpretation across them.

More broadly, our results suggest that resource-limited growth across biological systems shares a mathematical structure. The same trade-off principle appears to organize growth in nutrient-limited algae, bacteria allocating their proteome, *Daphnia* experiencing resource limitation, and human cancers exposed to cyclic hormone therapy. The trade-off exponent h is dimensionless and system-independent, allowing comparison across organisms and limitation regimes. The most direct cross-system application is in bacterial proteome allocation, where Scott-Hwa framework describes how ribosomal and metabolic fractions trade off under nutrient limitation (14), and recent extensions show that proteome reserves and growth-lag costs in fluctuating environments depend on how steeply fitness penalizes suboptimal allocation (58, 59), precisely what h quantifies. A natural extension is to multi-resource settings. Bacterial allocation strategies are already known to differ qualitatively across carbon, nitrogen, and phosphorus limitations (60), and stoichiometric frameworks couple multiple resources directly (12). In phytoplankton co-limited by nitrogen (N) and phosphorus (P), for example, a partition between N-uptake and P-processing machinery is structurally analogous, though whether the single-resource form transfers directly remains open. The canonical Redfield N:P ratio is one place where estimating h across taxa would carry direct interpretive weight (61, 62).

The framework is constrained by several key assumptions and by limitations in the data available to test them. The trade-off exponent is only inferred from population dynamics, so h remains mostly a phenomenological description of the system. We also exclude the analysis for h < 0; however, the generalized growth formulation itself allows for negative value of h. For example, for h = −1, G(q, Q, h) = 1/Q. This would imply growth in this regime depends only on the total budget and is wholly insensitive to how the resource is partitioned. Whether any real organism ever occupies this regime, and what biological structures would produce it, is an open question we do not pursue here. Both cohorts are monitored by PSA, an imperfect proxy for tumor burden, and because cancer mass is normalized initial value, the model’s prediction of cancer dynamics is only relative burden. On the statistical side, failure is a rare endpoint in the Bruchovsky cohort, so statistical test such as the Cox proportional-hazards model may be less robust and a larger cohort is needed to improve these results. Furthermore, the association between h and failure is only observational since we have not shown that h is modifiable, although it would be within our interpretation. The allocation of h is, however, directly testable. Single-cell and spatially resolved multi-omic profiling of serial biopsies through an androgen-deprivation cycle (63) should be able to resolve how resource pools are reorganized, by ionomic, metabolomic, or proteomic readout, as tumors shift from androgen-dependent growth toward alternative survival and proliferation programs. The framework makes the prediction that because h sets how steeply growth falls away from the optimum, low-h tumors should accommodate larger molecular reorganization for a smaller growth penalty than high-h tumors. Altogether, these results support h as a compact, dimensionless link between four seemingly different growth laws and a consistent indicator of the propensity of tumors to escape androgen deprivation across two cohorts, two drugs, and two disease states.

## Materials and Methods

**Intermittent androgen suppression therapy:** Longitudinal PSA and serum androgen data are from the Canadian intermittent androgen suppression trial (22), a Phase II study of intermittent ADT in men with biochemical recurrence after radiotherapy for locally advanced prostate cancer. Of 109 enrolled patients, 71 had complete data suitable (at least one complete cycle). **Adaptive abiraterone therapy:** Longitudinal PSA data from 32 men with metastatic castration-resistant prostate cancer enrolled in the first clinical trial of evolutionary-guided adaptive therapy (NCT02415621) (23). Serum androgen was not collected. Treatment failure was defined per the original trial protocol. Details of the cohorts are presented in the S1 text. The model was fit to PSA (and serum androgen when available) using a nonlinear mixed-effects framework (Monolix, Lixoft 2024), which estimates the population distribution and the individual parameters jointly. Details of the procedure are presented in the S5 Text.

## Data and Code Availability

All codes will be made available upon acceptance and prior to publication. Original cohort data are publicly available from the trial repositories.

## Acknowledgements

This research received no specific funding.

## Supporting Information for

### S1. Clinical Cohort and data

#### The Canadian Clinical Trial cohort for Intermittent Androgen Deprivation Therapy

(22) consists of 71 men who had complete longitudinal data suitable for nonlinear mixed-effects fitting. All patients had biochemical recurrence after radiotherapy for locally advanced prostate cancer, defined by the original trial protocol. The cohort comprises 6,114 longitudinal observations (3,057 each of PSA and serum androgen) spanning up to seven years of follow-up.

Each treatment cycle in the Canadian trial consisted of:

○ Four weeks of cyproterone acetate (CPA) lead-in therapy at 100 mg/day to prevent androgen flare on initiation of LHRH agonist therapy.
○ Combination leuprolide acetate plus continued CPA for the remainder of the ON period.
○ An OFF-treatment interval initiated when serum PSA dropped below a per-patient threshold and continued until PSA rose above a second threshold, at which point a new cycle began.

Treatment failure was defined as biochemical progression – a sustained rise in PSA during ON treatment indicating loss of androgen sensitivity – following the original trial protocol. Of the 71 patients in our analysis, 56 successfully completed the study without biochemical failure (success), and 15 experienced biochemical failure during follow-up (failure).

#### The Moffitt Adaptive Therapy cohort

(23) consists of 32 men with metastatic castration-resistant prostate cancer (mCRPC) enrolled in the first clinical trial of evolutionary-guided adaptive therapy (NCT02415621) who had complete longitudinal data suitable for nonlinear mixed-effects fitting. All patients had progressed on prior androgen deprivation therapy (i.e., PSA increases despite ADT) and met mCRPC criteria at trial entry, with continuous LHRH-agonist suppression (leuprolide) maintained as background therapy throughout follow-up in both arms. The cohort comprises 678 longitudinal PSA observations spanning up to 13 treatment cycles per patient (median 3) and up to 4.7 years of follow-up (median 1.8 years). Unlike the Canadian cohort, serum androgen was not collected longitudinally.

Patients were randomized between two arms:

Standard of care (SOC, n = 15): continuous abiraterone acetate at 1000 mg/day plus prednisone 5 mg twice daily, administered without interruption. Adaptive therapy (n = 17): identical dosing during ON periods, but with treatment cycled per individual PSA dynamics:

○ An ON period of abiraterone acetate (1000 mg/day) plus prednisone (5 mg twice daily).
○ An OFF interval initiated when serum PSA dropped to ≤50% of the pre-treatment baseline value.
○ A new ON period began when PSA returned to the pre-treatment baseline.

Treatment failure was defined as radiographic, symptomatic, or PSA progression per the original trial protocol, indicating loss of disease control under the assigned regimen. The Adaptive arm achieved a substantially longer time to progression than the SOC arm (median 916 vs 420 days). We remark on two points. First, while Zhang et al. mentioned 16 patients on SOC, the publicly available data associated with the publication only contains 15 patients. Secondly, these patients were also treated with an GnRH analog to suppress testicular androgen production. GnRH is not stopped during off-treatment in patients who were not surgically castrated. We do not have this information from the public data file.

### S2. Derivation of the generalized growth model

Consider the generalized growth function

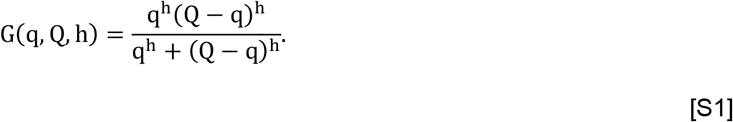

Assuming q, Q, h are positive, taking the partial derivative with respect to q gives:

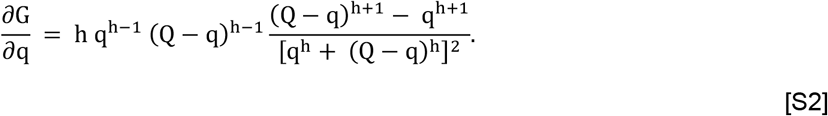

Solving 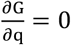 gives two roots, q = Q and 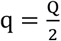. At q = Q, the growth function attains its minimum value G(q = Q) = 0, while at 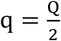, the growth function attains its maximum value 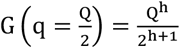. It is worth noting that for h < 1and q ≈ 0, the factor q^h−1^could lead to a steep selection gradient near the boundary.

To obtain Prater’s growth efficiency form, we observe that for 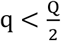 and large h, the growth function G(q, Q, h) approaches q^h^. On the other hand, for 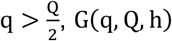 becomes (Q − q)^h^ in the large limit of h. Thus, we have that

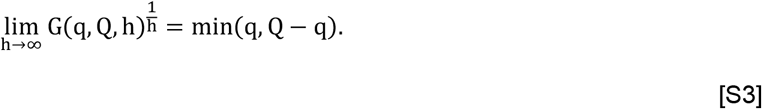

Recall that 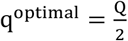, hence the difference from the optimal is defined as 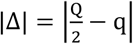 . This gives:

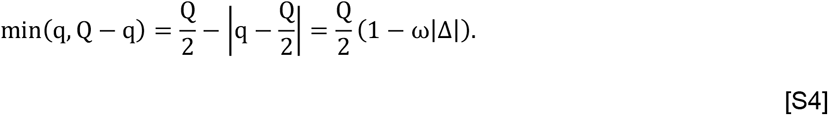

Where the scaling constant in Prater’s is taken to be 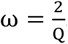. This derivation gives back Liebig’s limit of Prater’s growth efficiency form, and suggests the scaling is inversely proportional to the maximum allocation.

#### Derivation for a small allocation deviation from optimum

For a small deviation from optimal allocation 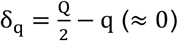, we have that 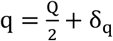 and 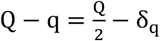. Substituting this to G(q, Q, h), we obtain:

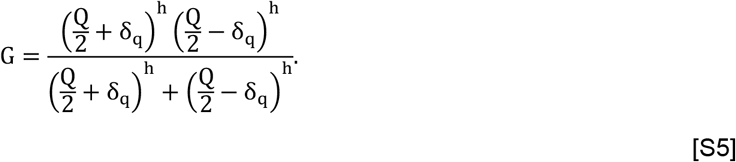

Factor 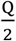, we obtain:

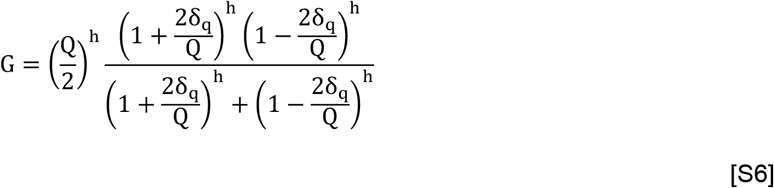

Let 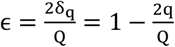 and simplify:

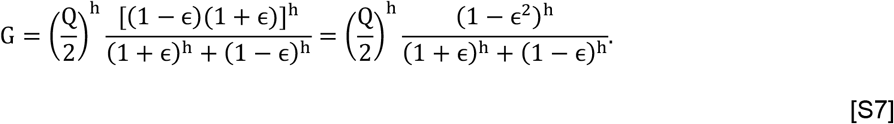

Now, Taylor expand with respect to small ϵ the numerator:

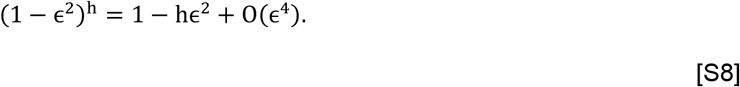

Similarly for the denominator, noting that 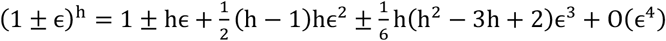, we obtain:

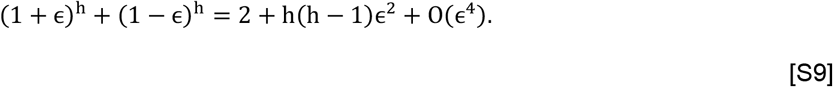

Putting these together and dropping the higher order term O(ϵ^4^):

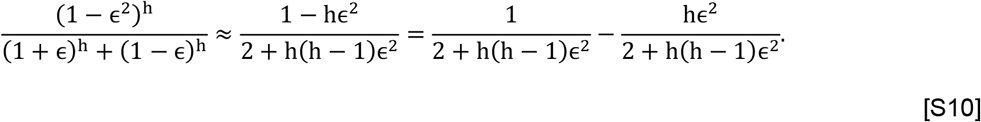

Note that the Taylor series for 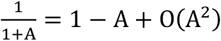, so using 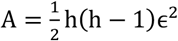, we obtain:

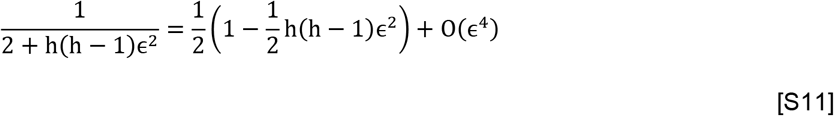

And

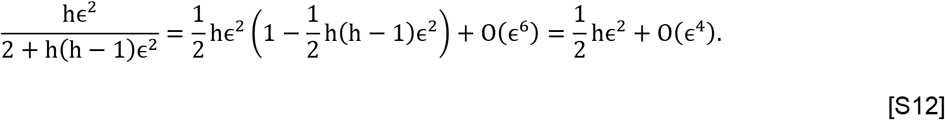

Dropping the higher order terms, we have:

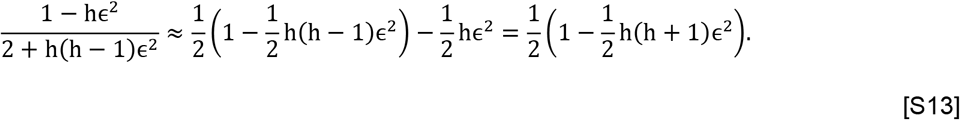

Therefore,

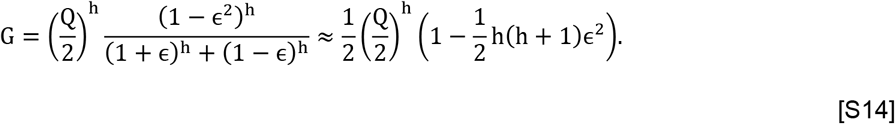

Substitute back 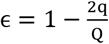, we get

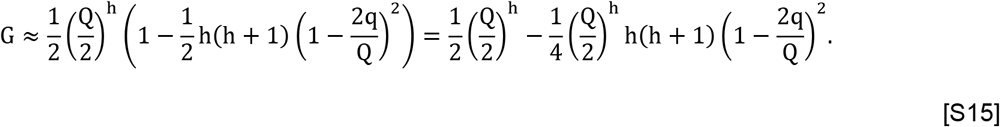

Note that the first term 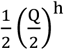 is the optimal growth at 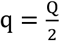, this means the small deviation from optimal allocation has a growth cost ΔG approximated by the second term:

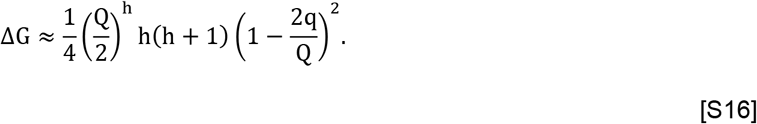

This term scales as h^2^ for large h. To arrive at this conclusion, one could alternatively start with Eq. S2 and find 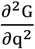 and evaluated at 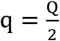.

### S3. A prostate cancer modeling framework

#### Breeder’s equation for the evolution of subsistence quota as population trait

We model the population-mean subsistence quota q as a quantitative trait evolving under selection. Lush’s breeder’s equation (28) gives the rate of change of a mean trait as the heritability times the selection gradient,

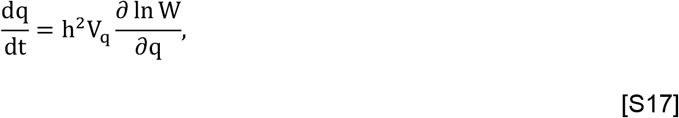

where h^2^ is heritability (a one-time separate notational use of the letter h from the trade-off exponent), V_q_ is trait variance, and W is fitness. Lumping heritability and variance into a single effective rate parameter ϵ_q_ and recognizing that the growth signal gradient is the log-fitness gradient, i.e., the log-fitness is the per-capita growth rate, (30) gives

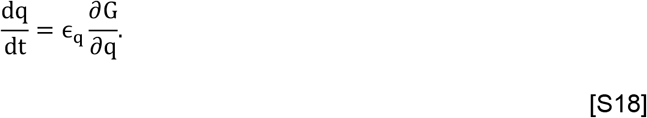

The selection gradient is positive when 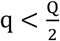 (selection pushes q up) and negative when 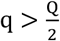 (selection pushes q down). A practical advantage of this formulation is that it does not require explicit tracking of separate androgen-sensitive and androgen-resistant subpopulations (inspired by Baez and Kuang(40)). Furthermore, we could have derived an equation for q(t) from the average of a sensitive and a resistant cancer subpopulation; however, such derivations are much more complicated and require additional assumptions on the trade-off between growth and resistance, such as the level of maximum resistance, and how much more fit the sensitive population is compared to the resistance population. Instead, the trade-off between sensitivity and resistance is already encoded in the shape of G(q, Q, h), making the single-population model with continuous subsistence quota trait suffices.

#### Cancer dynamics and evolution

Cancer growth dynamics follows net change equal growth minus death. The growth part follows from the generalized growth form μ_0_G(q, Q, h)x, while we use two death terms, a per capita death rate d_x_ and a density-dependent death rate δ. However, the maximum growth rate in a realistic system is likely not simply proportional to the subsistence cell quota, for example μ_m_ = μ_0_q, but likely depends on other factors unaccounted for in the generalized single resource growth law. Thus, we made the choice to decouple the maximum growth rate μ_m_ from its dependence on q. This gives us the equation governing cancer dynamics:

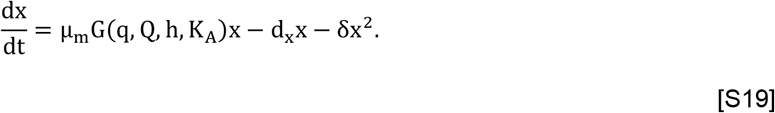

The function G(q, Q, h, K_A_ ) is the normalized G(q, Q, h) by multiplying with 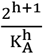. This normalizing constant follows from Eq. S2, where we show 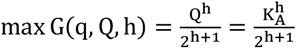, and K is the max cycle-1 androgen, rounded up to nearest 10 nmol/L. K_A_ represents the maximum androgen under no treatment. We initially considered taking the first serum androgen (prior to treatment) as its value. However, we note instances where androgen level recovered to higher than baseline. Many reasons could have caused this, since we only have a single serum androgen measurement, so it is less reliable to use it as the “highest” androgen level. In practice, this value could be taken directly from the patient’s health record. The explicit form of G(q, Q, h, K_A_) i_s_:

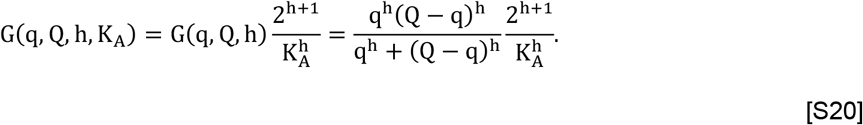

G captures only cancer proliferation (not net growth) (38, 40), so it is defined on the feasible domain 0 < q ≤ Q. Androgen suppression routinely drives Q below q during treatment, at which point the surplus Q − q is numerically fixed at a small positive floor (0.001 nmol/L), similar to previous work (38). The death terms d_x_ and δ are unaffected, and the population declines through turnover. Finally, we note that the coupling between the size of the cancer population and the PSA production introduces an identifiability issue. Thus, we made x the normalized cancer population with x(0) = 1.

#### Derivation of the intracellular androgen equation based on conservation law

The governing equation for intracellular androgen dQ/dt follows from conservation of total tumor androgen, in direct parallel with the derivation by Baez and Kuang (40) for their prostate cancer models. Consider Q(t) as the per-cell intracellular androgen concentration and x(t) as all cancer cell population. Then, the total androgen contained in the cancer population is Q_x_ = Qx. This is a conserved process involving the passive membrane diffusion into the cells, and the removal of androgen with dying cells. The inflow of androgen from the serum pool A to Q with rate proportional to the difference A − Q and to the available cellular volume x: v(A − Q)x, where v is the effective transport rate. The outflow from cell death which removes both intracellular androgen and the dying cells. The rate of cell loss in the model is (d_x_ + δx)x, so the outflow rate is (d_x_ + δx)Qx. The conservation law for total tumor androgen is therefore

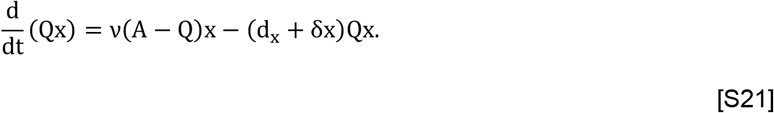

Applying the product rule to the left side and substituting dx/dt = (μ_m_G − d_x_ − δx)x:

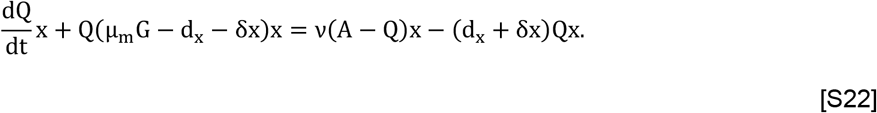

Dividing through by x and collecting like terms:

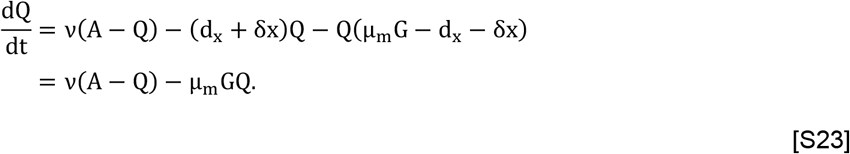

The cancer death terms cancel exactly. This cancellation is independent of the choice of the growth term. The first term is the net membrane flux from the serum pool (Fick-like diffusion). The second term is dilution by growth. As cells divide at rate μ_m_G, the per-cell quota Q is diluted because the total intracellular pool is shared among more cells.

The conservation argument was constructed in direct analogy with Droop’s nitrogen conservation for phytoplankton; however, it is not exact. In Droop’s setting, nitrogen is taken up actively, incorporated covalently into proteins and nucleic acids, and the per-cell quota is literally shared among daughter cells at division, with no pathway to rapidly re-equilibrate the quota with the environment. On the other hand, androgen is catalytic (binds androgen receptor, regulates transcription, then dissociates) rather than consumed, and the intracellular pool exchanges rapidly with serum via passive membrane diffusion. Nevertheless, the conservation law applies in both cases, since similar to the elements, androgen is not being changed in the process. Looking at the magnitude of the passive diffusion and dilution term, it is likely that passive diffusion dominates since μ_m_G is small to reflect tumor doubling time of week to months, while passive diffusion rate is on the order of hours to days. Thus, for simplicity, one could treat the dilution term as a negligible factor in most cases. We chose to keep it here as it maintains the structural conservation of the model.

#### Derivation for the production of prostate specific antigen (PSA)

Despite the name “prostate-specific antigen”, PSA is not uniquely produced by prostate cancer cells, or prostate cells in general. For modeling purposes, we often assume non-prostate source of PSA is negligible, which is an implicit assumption in the model. While healthy prostate cells also produce PSA, they are contained within the prostate epithelium wall. Cancerous growth can disrupt the epithelial barrier, which ultimately results in an increasing level of serum PSA in the blood via vascular leakiness (48), and often is the first sign of prostate cancer.

For PSA production by these cells, we first note that androgen receptor drives both PSA expression and proliferation (64), which inspired the idea of linking PSA production to prostate cell growth rate (38). For the PSA production by non-cancerous prostate cells, which we do not track in our model, we use serum androgen A level as a proxy for its proliferation: bA. Here, the constant b represents the effective PSA production rate of the changing non-cancerous cell population during IADT. When the treatment is on, serum androgen is suppressed, leading to slow growth of non-cancerous cells and lower PSA production. Note that since ADT does not completely shut off all androgen-producing sources, non-cancerous cells continue to produce a small amount of PSA during on-treatment. For the PSA production by cancer cells, we represent this using a hill function 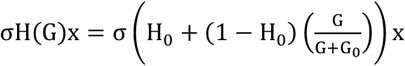 . Here, σ is the maximum PSA production rate, H(G) modifies the maximum PSA production rate based on the proliferation of cancer cells. H(G) ranges between H_0_ and 1 in our model to account for low level of PSA production under ADT. This also accounts for the possibility of a small subset of cancer that might not depend on androgen for growth (65, 66), such as the AR-V7 mutation (67). PSA is also assumed to clear at rate c. Altogether, we obtain the governing equation for PSA dynamics.

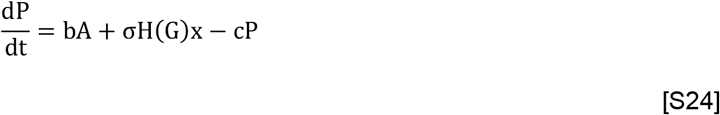

Now, let f_C_ be the fraction of cancer cell out of all PSA-producing cells at initial time. If we assume PSA is at a balance condition (for short-term) and set dP/dt = 0 at t = 0, we obtain:

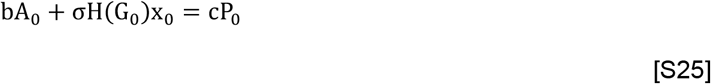

Expand both sides

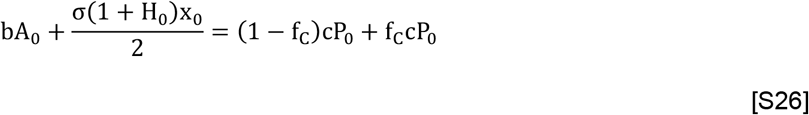

Note that the first term on each side of the equation corresponds to each other biologically since they both depict the fraction of PSA produced by non-cancerous cells. Similarly, the second terms correspond to the fraction of PSA produced by cancer cells at the initial time. This gives:

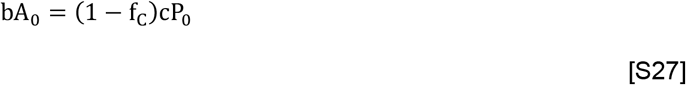

and

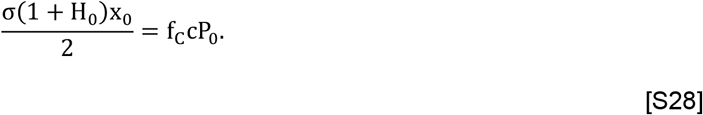

By solving both equations, we obtain closed forms approximation of parameters b and σ in term of the initial measurements:

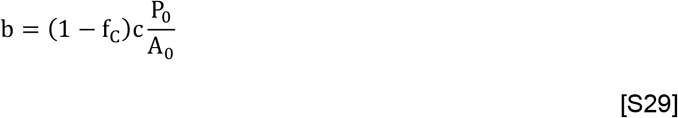

and

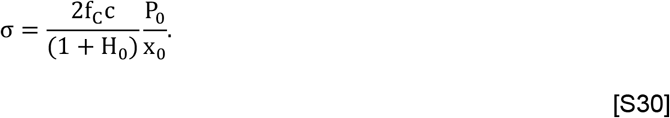

Here, 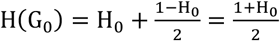 and x0 is scaled to 1.

#### Derivation of serum androgen dynamics

The framework so far has described how a cell’s growth depends on its internal allocation. In any realistic application, the environment itself is dynamic: the resource pool may be used up by the cells, by exogenous depletion, or by a treatment that perturbs supply. Thus, serum androgen also plays an important role in modulating the growth and evolution of cancer cells.

We model the production of serum androgen using a first-order saturation equation with production capacity K_A_ and production rate γ_e_, where it is the sum of the testes and adrenal gland production (γ_1_and γ_2_, respectively). We further fix γ_2_ to be 0.01 of γ_1_. Serum androgen decays at per capita rate d_A_ and is transferred between the serum and intracellular compartments at a rate proportional to the concentration difference A − Q (using the same assumption as in the derivation for intracellular androgen dynamics). The corresponding intracellular androgen gain is v(A − Q), whereas the serum loss is 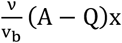 across cellular volume x, where v_b_ is the blood volume and Q ≤ A always holds since the intracellular androgen concentration is less than the serum concentration in this formulation. Together the equation governing serum androgen dynamics is:

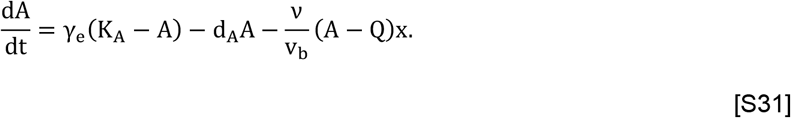

The treatment regime, as carried out in the clinical trial (22), only affect the testes production, hence we make γ_e_ a function of the effective treatment variable D, γ_e_(D) = (1− D)γ_1_+ γ_2_. The variable D goes from 0 (no suppression effect) to 1 (full suppression of testes production) as described by

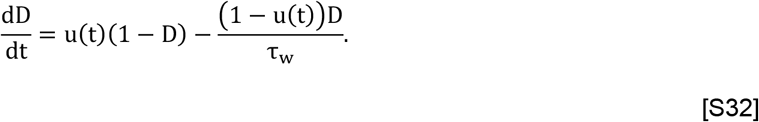

Here, u(t) is the treatment indicator. When treatment is on u(t) = 1, and dD/dt = 1− D, which rises up to 1. When the treatment is off u(t) = 0, and dD = −D/τ_w_, which decays to 0 with τ_w_ being the expected time for the drug effectiveness to go away.

In the IADT application, the clinical literature documents a progressive loss of androgen recovery across cycles. Tunn et al. (41) noted that across 109 patients, ∼80% reached pre-treatment androgen level during cycle 1, while ∼65% do during cycle 2. Pedraza and Kwart (42) noted four patients aged >70 with a median ADT of 108 months did not recover androgen level during the 3 years follow-up. In the

Bruchovsky data, we also observe a progressive loss of maximum androgen level from ∼75% recovery in cycle 1, to ∼50% in cycle 2, and so on. There has been just one previous model that accounts for this loss of androgen production (45). However, Reckell et al. model is rather complicated since it also accounts for the specific drug dynamics and interactions. A simpler model that accounts for this progressive loss is

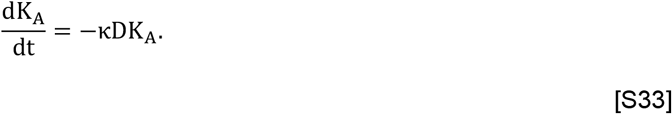

This turns the homeostatic serum androgen level into a dynamic variable that changes with treatment at effective rate κD.

#### The full prostate cancer model

The full system is present below:

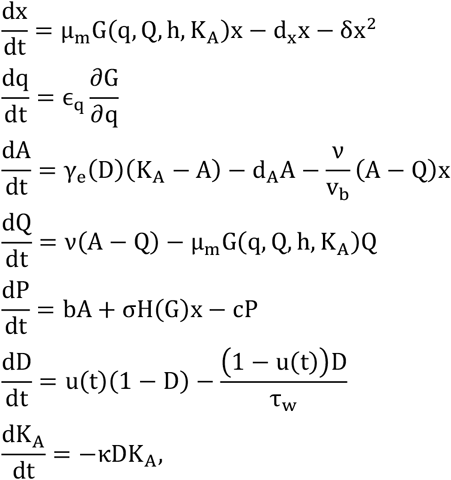

where

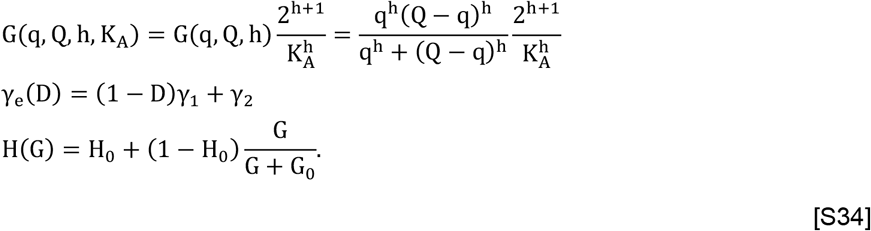

For the initial conditions of the system, we use x(0) = 1, q(0) = q_F_A_0_ (q_F_ is the initial quota fraction), A(0) = A_0_ (taken from the first observed serum androgen), Q(0) = A_0_ (rationale – serum at equilibrium), P(0) = P_0_ (first observed PSA), D(0) = 0 (no drug prior to treatment), K_A_(0) is the max cycle-1 androgen, rounded up to nearest 10 nmol/L.

### S4. A simplified prostate cancer applied to the adaptive therapy cohort

The publicly available data for the adaptive therapy cohort did not contain androgen observation (23), which does not allow for direct application of the full prostate cancer model. Thus, we make several strong assumptions to simplify the model appropriately.

We remove the androgen variables: A, Q, K_A_. The effect of the drug (abiraterone) is incorporated by a normalized variable A_e_ = 1− D, where D is the drug variable (same as before). We then let the effective intracellular androgen Q_e_ = A_e_. Since A_e_ is normalized, we now estimate q_0_ directly (instead of q_F_). The simplified system becomes:

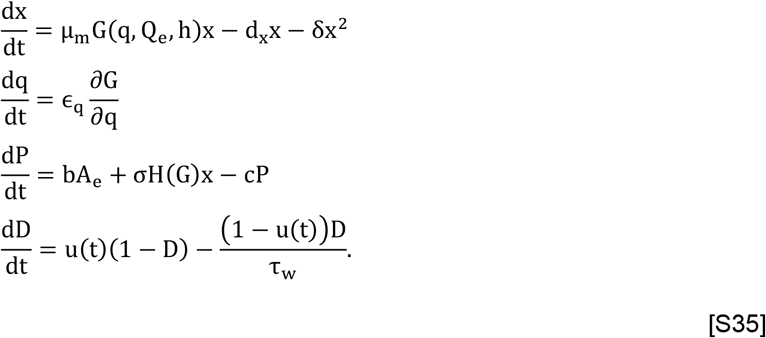

We remark that this formulation is consistent with the mechanism of action for abiraterone, which is an androgen biosynthesis inhibitor that blocks CYP17A1 enzyme required for androgen synthesis (68). Hence, it suppresses androgen production in the testes, adrenal glands, and cancer cells. A model from the same cell-quota family was previous applied to a cohort treated with abiraterone at Mayo Clinic (69). Besides abiraterone, patients in the adaptive cohort also receive a gonadotropin-releasing hormone (GnRH) analog treatment to suppress testes androgen production. Patients who did not undergo surgical castration remains on GnRH analog even during the off-treatment period. This likely affects the estimate of τ_w_ for this cohort and the precise interpretation of A_e_ and Q_e_.

### S5. Numerical methods

#### Population fitting with nonlinear mixed effect framework

We used the Stochastic Approximation Expectation-Maximization (SAEM) algorithm implemented in Monolix (Lixoft, 2024). We minimize the error in PSA and androgen, assuming a constant error model. In total, we fit 10 model parameters with random effect (μ_m_, h, ϵ_q_, γ_1_, d_A_, f_C_, d_x_, κ, q_F_, δ), and 2 model parameters without random effect (τ_w_, H_0_). All parameters are fitted using lognormal distribution, except for f_C_ and q_F_ which are logit normal in the feasible range (0,1). Group comparison between success (n = 56) and failure (n = 15) used the two-sided Mann-Whitney U-test. We also fit the simplified model to the adaptive therapy cohort. In total, we have 7 parameters with random effects (μ_m_, h, ϵ_q_, f_C_, d_x_, q_F_, δ) under the same setting. Model fit and parameter stratification for the adaptive cohort is presented in Fig. S3. A demonstration of model agreement with the data using *R*^2^ is shown in Fig. S4 (Bruchovsky cohort) and S5 (Moffitt cohort).

#### Identifiability of the trade-off exponent h and other model parameters

All model estimated parameters appear to be well-identified by Monolix: relative standard errors (RSE) on population parameters are within 25% (Table S2). In particular, the RSE for h is just 6.4%. The random effect parameters are also indicated to be well-quantified with clearly bounded range (Table S3).

Parameters for the simplified model fit to the adaptive therapy cohort is less well-quantified likely because the androgen observations are not. These results are presented in Table S4 (population estimates) and Table S5 (individual estimates).

#### Survival analysis on individual model parameters

We tested whether the patient-specific parameters from the full prostate cancer model could predict which patients would fail treatment. Time to treatment failure came from the clinical records (22). We fit a Cox model using the 10 parameters that vary between patients as predictors. The model assigns each patient a risk score, and we split patients at the median score into a low-risk group (36 patients) and a high-risk group (35 patients). We tested the model two ways. The in-sample version (Fig. 4A) fits on all 71 patients and scores the same patients. The leave-one-out version (Fig. 4B) fits the model 71 times, each time leaving out one patient and scoring that patient with a model that fit only the remaining 70 patients. We report three statistics: the hazard ratio (HR) between the two groups with 95% Wald confidence interval, estimated from a Cox regression with the binary risk-group indicator as the sole predictor; the log-rank p-value for the difference between the two Kaplan–Meier survival curves; and Harrell’s concordance index C computed from the underlying multivariate Cox model’s linear predictor. Kaplan–Meier curves are plotted with 95% Greenwood pointwise confidence bands. Vertical ticks indicate censoring times. All analyses follow standard methodology (70).

## Tables

**Table S1.**
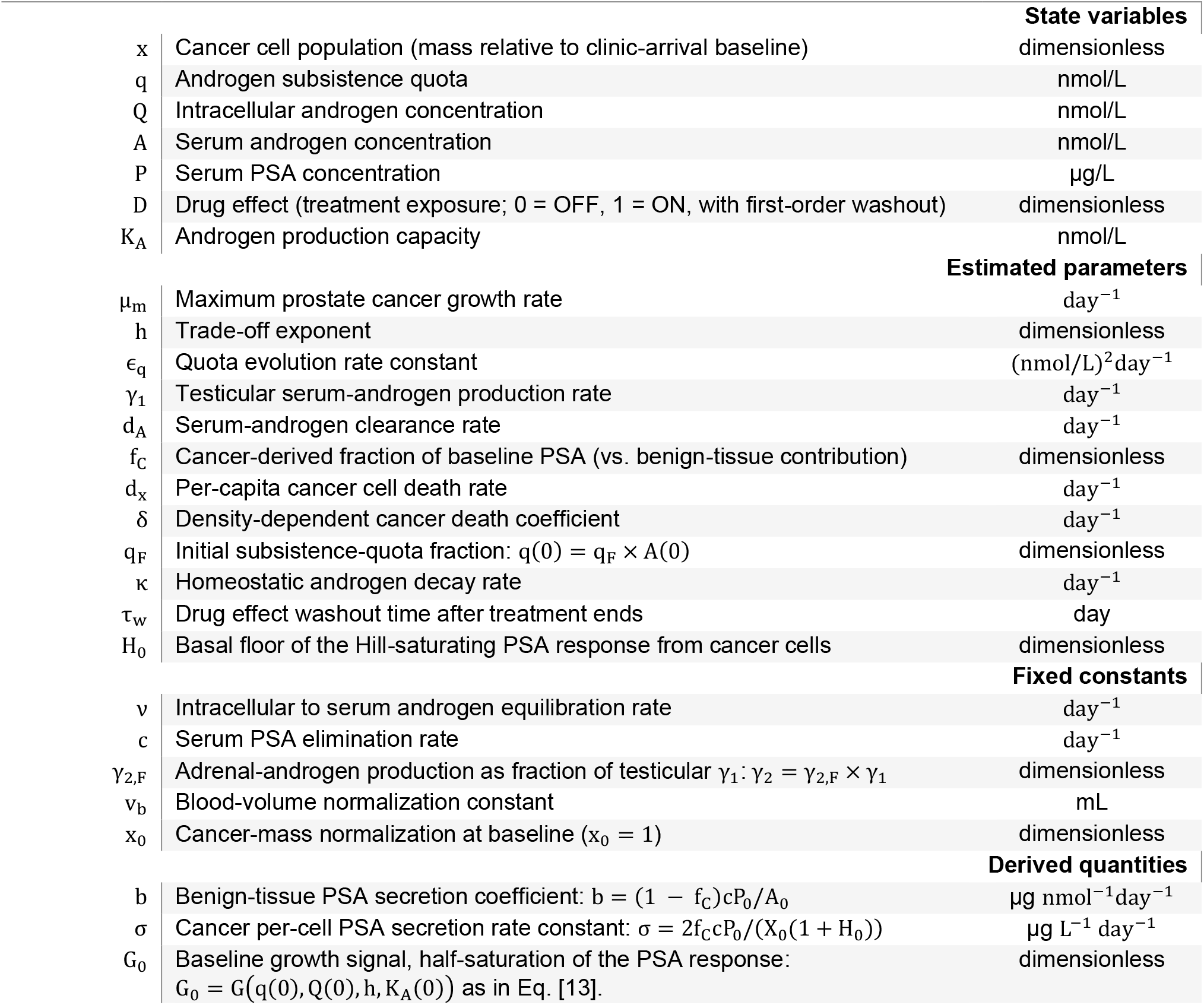
Model parameters, state variables, regressors, and derived quantities.

**Table S2.** Population parameter priors and fitted estimates (Bruchovsky cohort).

| Parameter | Value | S.E. | R.S.E. (%) | C.V. (%) | 2.5% CI | 97.5% CI |
| --- | --- | --- | --- | --- | --- | --- |
| **Fixed effects** |  |  |  |  |  |  |
| $\mu_m$ | 0.052 | 0.0047 | 9.06 | — | 0.044 | 0.062 |
| $h$ | 0.61 | 0.039 | 6.38 | — | 0.54 | 0.69 |
| $\epsilon_q$ | 0.0018 | 3.70e-04 | 20.3 | — | 0.0012 | 0.0027 |
| $\gamma_1$ | 0.33 | 0.049 | 15 | — | 0.24 | 0.43 |
| $d_A$ | 0.099 | 0.0098 | 9.91 | — | 0.082 | 0.12 |
| $v$ | 1 | — | — | — | — | — |
| $f_C$ | 0.98 | 0.019 | 1.97 | — | 0.92 | 0.99 |
| $d_X$ | 0.0039 | 5.90e-04 | 14.9 | — | 0.003 | 0.0053 |
| $\kappa$ | 7.30e-04 | 7.90e-05 | 10.8 | — | 5.90e-04 | 9.00e-04 |
| $q_F$ | 0.63 | 0.075 | 12.1 | — | 0.47 | 0.76 |
| $\tau_w$ | 31.2 | 0.065 | 0.207 | — | 31.1 | 31.3 |
| $c$ | 0.25 | — | — | — | — | — |
| $\delta$ | 0.013 | 0.0031 | 24.2 | — | 0.008 | 0.02 |
| $\gamma_{2,F}$ | 0.01 | — | — | — | — | — |
| $v_b$ | 5000 | — | — | — | — | — |
| $H_0$ | 0.15 | 0.0028 | 1.84 | — | 0.15 | 0.16 |
| **Standard deviation of the random effects** |  |  |  |  |  |  |
| $\omega_{\mu_m}$ | 0.72 | 0.068 | 9.44 | 82.4 | 0.6 | 0.87 |
| $\omega_h$ | 0.49 | 0.055 | 11.2 | 52 | 0.39 | 0.61 |
| $\omega_{\epsilon_q}$ | 1.43 | 0.16 | 11.5 | 259 | 1.14 | 1.78 |
| $\omega_{\gamma_1}$ | 1.17 | 0.12 | 10 | 171 | 0.96 | 1.42 |
| $\omega_{d_A}$ | 0.76 | 0.067 | 8.86 | 88.5 | 0.64 | 0.9 |
| $\omega_{f_C}$ | 3.39 | 0.7 | 20.6 | 30.9 | 2.29 | 5.01 |
| $\omega_{d_X}$ | 0.99 | 0.12 | 12.1 | 130 | 0.79 | 1.26 |
| $\omega_{\kappa}$ | 0.81 | 0.076 | 9.4 | 96.7 | 0.68 | 0.98 |
| $\omega_{q_F}$ | 2.36 | 0.23 | 9.68 | 58.4 | 1.95 | 2.85 |
| $\omega_{\delta}$ | 1.9 | 0.17 | 9.14 | 604 | 1.59 | 2.27 |
| **Error model parameters** |  |  |  |  |  |  |
| $a$ | 1.84 | 0.026 | 1.42 | — | 1.79 | 1.89 |
| $a_2$ | 2.3 | 0.03 | 1.3 | — | 2.24 | 2.36 |

**Table S3.** Individual parameter estimates (Bruchovsky cohort).

| ID | $\mu_m$ | $h$ | $\epsilon_q$ | $\gamma_1$ | $d_A$ | $f_C$ | $d_X$ | $\kappa$ | $q_F$ | $\delta$ |
| --- | --- | --- | --- | --- | --- | --- | --- | --- | --- | --- |
| 1 | 0.0243 | 0.38 | 2.66e-04 | 0.0944 | 0.0982 | 0.998 | 0.00104 | 6.05e-04 | 0.526 | 0.0246 |
| 2 | 0.0819 | 0.505 | 0.00301 | 0.0786 | 0.0771 | 0.928 | 0.00466 | 6.69e-04 | 0.722 | 0.0378 |
| 3 | 0.0922 | 0.722 | 0.0181 | 0.132 | 0.0303 | 0.998 | 0.00245 | 5.29e-04 | 0.714 | 0.0583 |
| 6 | 0.102 | 1.45 | 0.00193 | 0.0602 | 0.0316 | 0.998 | 0.00958 | 8.96e-04 | 0.106 | 0.0539 |
| 7 | 0.116 | 0.207 | 4.97e-04 | 0.917 | 0.178 | 0.987 | 0.00332 | 7.43e-04 | 0.902 | 0.165 |
| 8 | 0.372 | 0.342 | 6.57e-04 | 0.145 | 0.154 | 0.996 | 0.0026 | 0.00118 | 0.975 | 0.271 |
| 12 | 0.136 | 0.941 | 3.70e-04 | 0.972 | 0.0667 | 0.992 | 0.00376 | 0.00223 | 0.0409 | 0.0792 |
| 13 | 0.17 | 0.661 | 0.0172 | 0.34 | 0.177 | 0.997 | 0.00352 | 2.67e-04 | 0.557 | 0.172 |
| 14 | 0.0298 | 0.81 | 0.00181 | 0.369 | 0.0797 | 0.996 | 0.00678 | 3.03e-04 | 0.378 | 0.0149 |
| 15 | 0.0433 | 0.48 | 0.0473 | 0.0843 | 0.127 | 0.977 | 0.00143 | 3.29e-04 | 0.518 | 0.0563 |
| 16 | 0.0782 | 0.747 | 0.00292 | 0.897 | 0.126 | 0.995 | 0.00407 | 6.87e-04 | 0.357 | 6.20e-04 |
| 17 | 0.0546 | 1.04 | 9.15e-04 | 0.577 | 0.154 | 0.994 | 0.00429 | 5.06e-04 | 0.116 | 0.0257 |
| 19 | 0.29 | 0.385 | 7.43e-04 | 1.05 | 0.172 | 0.995 | 0.00327 | 0.00237 | 0.986 | 0.489 |
| 22 | 0.021 | 0.469 | 0.00185 | 0.0627 | 0.155 | 0.991 | 0.00676 | 8.85e-04 | 0.538 | 0.00368 |
| 24 | 0.079 | 0.703 | 4.34e-04 | 0.429 | 0.0244 | 0.993 | 0.00153 | 9.08e-04 | 0.357 | 0.133 |
| 25 | 0.0538 | 0.916 | 3.28e-04 | 1.01 | 0.0709 | 0.0879 | 0.00275 | 0.00152 | 0.00786 | 2.25e-04 |
| 26 | 0.0788 | 0.969 | 7.83e-04 | 1.28 | 0.0475 | 0.975 | 0.00188 | 0.0012 | 0.227 | 0.133 |
| 28 | 0.0177 | 0.892 | 5.97e-04 | 0.623 | 0.173 | 0.999 | 0.00319 | 6.81e-04 | 0.221 | 0.0289 |
| 29 | 0.0199 | 0.917 | 0.00292 | 0.821 | 0.155 | 0.951 | 0.00823 | 8.83e-04 | 0.36 | 0.00522 |
| 30 | 0.0251 | 0.741 | 0.00212 | 1.41 | 0.064 | 0.988 | 0.00352 | 0.00149 | 0.653 | 0.0303 |
| 31 | 0.0415 | 0.622 | 0.00363 | 0.0556 | 0.139 | 0.995 | 0.00527 | 0.00217 | 0.605 | 0.0194 |
| 32 | 0.0671 | 0.469 | 6.36e-04 | 0.115 | 0.0215 | 0.614 | 0.0257 | 5.27e-04 | 0.151 | 0.00209 |
| 33 | 0.0152 | 0.298 | 7.02e-04 | 0.0631 | 0.144 | 0.0656 | 0.00131 | 4.76e-04 | 0.242 | 2.93e-04 |
| 36 | 0.105 | 0.356 | 9.70e-04 | 0.271 | 0.11 | 0.995 | 0.0165 | 4.69e-04 | 0.514 | 0.0267 |
| 37 | 0.0986 | 0.474 | 0.00115 | 0.143 | 0.259 | 0.646 | 0.00157 | 8.61e-04 | 0.582 | 0.0304 |
| 39 | 0.0301 | 0.322 | 1.81e-04 | 0.142 | 0.126 | 0.982 | 0.00152 | 6.51e-04 | 0.579 | 0.0242 |
| 40 | 0.0396 | 0.755 | 0.00179 | 0.151 | 0.194 | 0.998 | 0.00715 | 4.19e-04 | 0.818 | 0.013 |
| 41 | 0.0237 | 0.428 | 4.14e-04 | 0.318 | 0.137 | 0.997 | 0.00619 | 8.24e-04 | 0.382 | 0.0105 |
| 42 | 0.0402 | 0.551 | 0.00186 | 0.216 | 0.0952 | 0.988 | 0.00337 | 0.00228 | 0.374 | 0.0222 |
| 44 | 0.0226 | 0.621 | 6.82e-04 | 0.517 | 0.0414 | 0.991 | 0.00583 | 0.00433 | 0.364 | 0.0177 |
| 46 | 0.0653 | 0.421 | 0.00119 | 0.135 | 0.0237 | 0.998 | 0.0305 | 0.00295 | 0.944 | 8.58e-04 |
| 48 | 0.0265 | 0.378 | 0.00136 | 0.364 | 0.143 | 0.99 | 0.00222 | 6.89e-04 | 0.701 | 0.00222 |
| 50 | 0.0586 | 0.204 | 5.61e-04 | 1.25 | 0.238 | 0.957 | 0.032 | 7.58e-04 | 0.999 | 0.018 |
| 51 | 0.0551 | 0.432 | 0.00447 | 0.952 | 0.0668 | 0.995 | 0.0036 | 0.00143 | 0.827 | 0.0386 |
| 52 | 0.109 | 0.309 | 7.38e-04 | 2.28 | 0.222 | 0.716 | 0.0233 | 7.42e-04 | 0.921 | 0.0223 |
| 54 | 0.0619 | 0.408 | 0.00179 | 0.0659 | 0.0427 | 0.357 | 0.0105 | 2.35e-04 | 0.801 | 9.48e-04 |
| 55 | 0.045 | 1.6 | 0.106 | 0.246 | 0.0541 | 0.536 | 0.00384 | 2.39e-04 | 0.985 | 7.52e-04 |
| 58 | 0.0527 | 1.46 | 0.00155 | 0.292 | 0.0467 | 0.995 | 0.00298 | 7.14e-04 | 0.353 | 0.0575 |
| 60 | 0.0735 | 0.529 | 9.26e-04 | 0.126 | 0.119 | 0.752 | 0.0151 | 5.14e-04 | 0.733 | 0.00629 |
| 61 | 0.0362 | 0.533 | 0.0011 | 0.225 | 0.189 | 0.979 | 0.00185 | 4.49e-04 | 0.119 | 0.00576 |
| 62 | 0.0278 | 0.709 | 6.67e-04 | 1.35 | 0.102 | 0.989 | 0.00408 | 0.00111 | 0.383 | 0.0344 |
| 63 | 0.0665 | 0.531 | 0.00235 | 0.169 | 0.0885 | 0.581 | 0.00284 | 3.36e-04 | 0.698 | 0.0201 |
| 64 | 0.0946 | 0.467 | 6.52e-04 | 0.598 | 0.0729 | 0.0226 | 0.00662 | 0.0032 | 0.983 | 3.41e-05 |
| 66 | 0.0507 | 0.974 | 0.00237 | 0.33 | 0.169 | 0.992 | 0.00652 | 2.81e-04 | 0.145 | 0.0108 |
| 71 | 0.065 | 0.54 | 0.0014 | 0.301 | 0.133 | 0.991 | 0.00294 | 0.00111 | 0.513 | 0.0154 |
| 75 | 0.0913 | 1.62 | 0.00825 | 0.396 | 0.113 | 0.997 | 0.00238 | 5.29e-04 | 0.192 | 0.00324 |
| 77 | 0.0575 | 0.822 | 5.36e-04 | 1.01 | 0.0816 | 0.987 | 0.00109 | 7.41e-04 | 0.376 | 0.0508 |
| 78 | 0.0559 | 0.381 | 3.08e-04 | 0.112 | 0.0947 | 0.736 | 0.0043 | 2.98e-04 | 0.967 | 0.00937 |
| 79 | 0.0547 | 0.384 | 4.63e-04 | 0.36 | 0.0445 | 0.852 | 0.00533 | 4.25e-04 | 0.698 | 0.0529 |
| 83 | 0.0483 | 0.469 | 0.00765 | 1.63 | 0.095 | 0.997 | 0.0025 | 8.49e-04 | 0.78 | 0.0602 |
| 84 | 0.062 | 0.522 | 0.00395 | 0.146 | 0.178 | 0.892 | 0.00353 | 0.00158 | 0.531 | 0.031 |
| 85 | 0.0244 | 0.351 | 7.79e-04 | 0.445 | 0.162 | 0.997 | 0.00794 | 0.00146 | 0.659 | 1.97e-04 |
| 86 | 0.0196 | 0.514 | 0.00137 | 0.0927 | 0.192 | 0.99 | 0.00247 | 9.92e-04 | 0.535 | 0.00679 |
| 87 | 0.0461 | 0.606 | 5.64e-04 | 0.117 | 0.0262 | 0.996 | 0.00126 | 0.00132 | 0.281 | 0.0539 |
| 88 | 0.0479 | 0.51 | 0.00328 | 0.222 | 0.0678 | 0.992 | 0.0036 | 9.73e-04 | 0.807 | 0.0224 |
| 91 | 0.0724 | 0.704 | 0.00432 | 0.268 | 0.164 | 0.642 | 0.00245 | 9.52e-05 | 0.928 | 0.0392 |
| 93 | 0.0319 | 1.04 | 0.0038 | 0.183 | 0.0695 | 0.994 | 0.00244 | 3.92e-04 | 0.504 | 0.0277 |
| 94 | 0.0374 | 0.757 | 0.00157 | 0.247 | 0.0895 | 0.998 | 0.00612 | 0.00165 | 0.854 | 0.0272 |
| 95 | 0.0221 | 0.932 | 0.0048 | 0.336 | 0.0452 | 0.96 | 0.00291 | 3.54e-04 | 0.722 | 0.0029 |
| 96 | 0.0386 | 0.759 | 5.65e-04 | 0.247 | 0.0828 | 0.99 | 0.00651 | 0.00153 | 0.801 | 0.023 |
| 97 | 0.0298 | 0.38 | 0.00231 | 0.0695 | 0.0421 | 0.981 | 0.00863 | 8.33e-04 | 0.647 | 0.0119 |
| 99 | 0.0389 | 0.759 | 0.0036 | 4.2 | 0.0772 | 0.999 | 0.0126 | 0.00345 | 0.932 | 0.00122 |
| 100 | 0.0523 | 1.02 | 0.00302 | 0.811 | 0.0798 | 0.994 | 0.00712 | 6.47e-04 | 0.724 | 0.0309 |
| 101 | 0.0674 | 0.65 | 0.002 | 1.42 | 0.379 | 0.997 | 0.00533 | 0.0021 | 0.935 | 1.11e-04 |
| 102 | 0.0406 | 0.78 | 0.00404 | 0.633 | 0.0476 | 0.843 | 0.00287 | 0.00133 | 0.779 | 0.00443 |
| 104 | 0.0399 | 0.456 | 0.00393 | 1.08 | 0.154 | 0.99 | 0.0135 | 5.87e-04 | 0.896 | 0.00807 |
| 105 | 0.0727 | 0.423 | 0.00197 | 0.441 | 0.138 | 0.996 | 0.0131 | 0.00112 | 0.884 | 0.0216 |
| 106 | 0.0183 | 0.852 | 0.00421 | 0.502 | 0.119 | 0.99 | 0.00473 | 0.00128 | 0.333 | 0.00841 |
| 107 | 0.0447 | 0.743 | 0.00229 | 0.374 | 0.107 | 0.98 | 0.00547 | 8.90e-04 | 0.567 | 0.0222 |
| 108 | 0.018 | 0.696 | 0.00905 | 0.747 | 0.0936 | 0.994 | 0.00332 | 4.34e-04 | 0.492 | 0.0131 |
| 109 | 0.0391 | 0.586 | 0.00152 | 0.484 | 0.076 | 0.985 | 0.00335 | 0.00261 | 0.518 | 0.0394 |

**Table S4.** Population parameter priors and fitted estimates (adaptive therapy cohort).

| Parameter | Value | S.E. | R.S.E. (%) | C.V. (%) | 2.5% CI | 97.5% CI |
| --- | --- | --- | --- | --- | --- | --- |
| Fixed effects |  |  |  |  |  |  |
| $\mu_m$ | 0.016 | 0.0074 | 44.9 | — | 0.0075 | 0.036 |
| $h$ | 0.47 | 0.053 | 11.1 | — | 0.38 | 0.59 |
| $\epsilon_q$ | 1.70e-05 | 2.60e-04 | **1530** | — | 6.50e-07 | 4.40e-04 |
| $f_c$ | 0.86 | 0.26 | 30.6 | — | 0.1 | 0.99 |
| $d_x$ | 8.70e-05 | 1.45e+04 | **1.67e+10** | — | 1.70e-08 | 0.44 |
| $q_F$ | 0.91 | 0.23 | 24.8 | — | 0.18 | 0.99 |
| $\delta$ | 0 | ∞ | ∞ | — | 0 | 1.40e-08 |
| $\tau_w$ | 120 | NaN | NaN | — | NaN | NaN |
| $c$ | 0.25 | — | — | — | — | — |
| $h_0$ | 0.1 | — | — | — | — | — |
| Standard deviation of the random effects |  |  |  |  |  |  |
| $\omega_{\mu_m}$ | 0.5 | 0.49 | **96.5** | **53.8** | 0.13 | 1.91 |
| $\omega_h$ | 0.4 | 0.19 | 46.8 | 41.8 | 0.18 | 0.9 |
| $\omega_{\epsilon_q}$ | 2.76 | 3.22 | **117** | **4521** | 0.63 | 12.1 |
| $\omega_{f_c}$ | 2.82 | 1.4 | 49.6 | 45.6 | 1.21 | 6.56 |
| $\omega_{d_x}$ | 2.86 | 2.9 | **102** | **5964** | 0.73 | 11.3 |
| $\omega_{q_F}$ | 2.18 | 0.91 | 41.7 | 31.1 | 1.04 | 4.55 |
| $\omega_{\delta}$ | 13.2 | 11.8 | **89.5** | ∞ | 3.7 | 47.3 |
| Error model parameters |  |  |  |  |  |  |
| $a$ | 0.021 | 0.03 | **143** | — | 0.0041 | 0.11 |
| $b_{psa}$ | 0.45 | 0.02 | 4.5 | — | 0.41 | 0.49 |

**Table S5.** Individual parameter estimates (adaptive therapy cohort). All individual estimates for δ converges to 0 (non-identifiable). SOC stands for standard-of-care.

| ID | Arm | $\mu_m$ | h | $\epsilon_q$ | $f_c$ | $d_x$ | $q_F$ |
| --- | --- | --- | --- | --- | --- | --- | --- |
| 1001 | Adaptive | 0.0291 | 0.843 | 0.00154 | 0.158 | 2.89e-05 | 0.947 |
| 1002 | Adaptive | 0.011 | 0.391 | 3.50e-05 | 0.888 | 5.80e-05 | 0.729 |
| 1003 | Adaptive | 0.0316 | 0.933 | 4.97e-06 | 0.993 | 0.00204 | 0.804 |
| 1004 | Adaptive | 0.013 | 0.416 | 1.24e-05 | 0.906 | 3.88e-05 | 0.986 |
| 1005 | Adaptive | 0.0173 | 0.376 | 1.43e-05 | 0.965 | 0.00217 | 0.678 |
| 1006 | Adaptive | 0.011 | 0.544 | 2.00e-04 | 0.0539 | 2.67e-05 | 0.685 |
| 1007 | Adaptive | 0.0347 | 0.306 | 1.81e-05 | 0.957 | 0.0224 | 0.757 |
| 1009 | Adaptive | 0.0177 | 0.475 | 6.15e-06 | 0.975 | 6.90e-04 | 0.934 |
| 1010 | Adaptive | 0.0185 | 0.368 | 1.56e-05 | 0.961 | 3.14e-05 | 0.97 |
| 1011 | Adaptive | 0.024 | 0.353 | 1.15e-05 | 0.127 | 0.00695 | 0.36 |
| 1012 | Adaptive | 0.00574 | 0.497 | 9.06e-05 | 0.57 | 0.00119 | 0.86 |
| 1014 | Adaptive | 0.013 | 0.516 | 8.79e-06 | 0.689 | 4.61e-05 | 0.902 |
| 1015 | Adaptive | 0.0113 | 0.433 | 1.38e-05 | 0.563 | 4.26e-05 | 0.474 |
| 1016 | Adaptive | 0.0182 | 0.429 | 3.01e-06 | 0.133 | 3.82e-05 | 0.959 |
| 1017 | Adaptive | 0.0265 | 0.927 | 3.38e-04 | 0.972 | 0.016 | 0.946 |
| 1018 | Adaptive | 0.016 | 0.407 | 1.74e-05 | 0.946 | 1.06e-04 | 0.987 |
| 1020 | Adaptive | 0.0162 | 0.431 | 8.10e-06 | 0.728 | 8.65e-05 | 0.96 |
| 2001 | SOC | 0.0156 | 0.424 | 6.99e-06 | 0.944 | 0.00207 | 0.982 |
| 2002 | SOC | 0.0149 | 0.544 | 2.97e-04 | 0.968 | 9.75e-04 | 0.975 |
| 2003 | SOC | 0.0147 | 0.452 | 9.48e-06 | 0.834 | 6.02e-05 | 0.965 |
| 2004 | SOC | 0.0209 | 0.347 | 1.60e-05 | 0.739 | 7.31e-05 | 0.897 |
| 2005 | SOC | 0.0289 | 0.418 | 7.20e-06 | 0.961 | 3.93e-05 | 0.766 |
| 2006 | SOC | 0.0188 | 0.338 | 7.23e-06 | 0.169 | 7.08e-05 | 0.922 |
| 2007 | SOC | 0.0176 | 0.36 | 1.06e-05 | 0.0887 | 4.45e-05 | 0.916 |
| 2008 | SOC | 0.0163 | 0.664 | 2.14e-05 | 0.61 | 2.25e-04 | 0.811 |
| 2009 | SOC | 0.014 | 0.606 | 1.45e-05 | 0.92 | 9.67e-05 | 0.876 |
| 2010 | SOC | 0.0145 | 0.48 | 9.40e-06 | 0.658 | 6.49e-05 | 0.974 |
| 2011 | SOC | 0.018 | 0.593 | 1.36e-05 | 0.964 | 7.77e-05 | 0.732 |
| 2012 | SOC | 0.0189 | 0.467 | 1.02e-05 | 0.147 | 7.49e-05 | 0.876 |
| 2013 | SOC | 0.0134 | 0.513 | 7.21e-06 | 0.929 | 7.06e-05 | 0.972 |
| 2014 | SOC | 0.0147 | 0.5 | 1.36e-05 | 0.924 | 2.61e-05 | 0.741 |
| 2015 | SOC | 0.017 | 0.373 | 1.64e-05 | 0.933 | 8.25e-05 | 0.963 |

## Figures

**Fig. S1.**
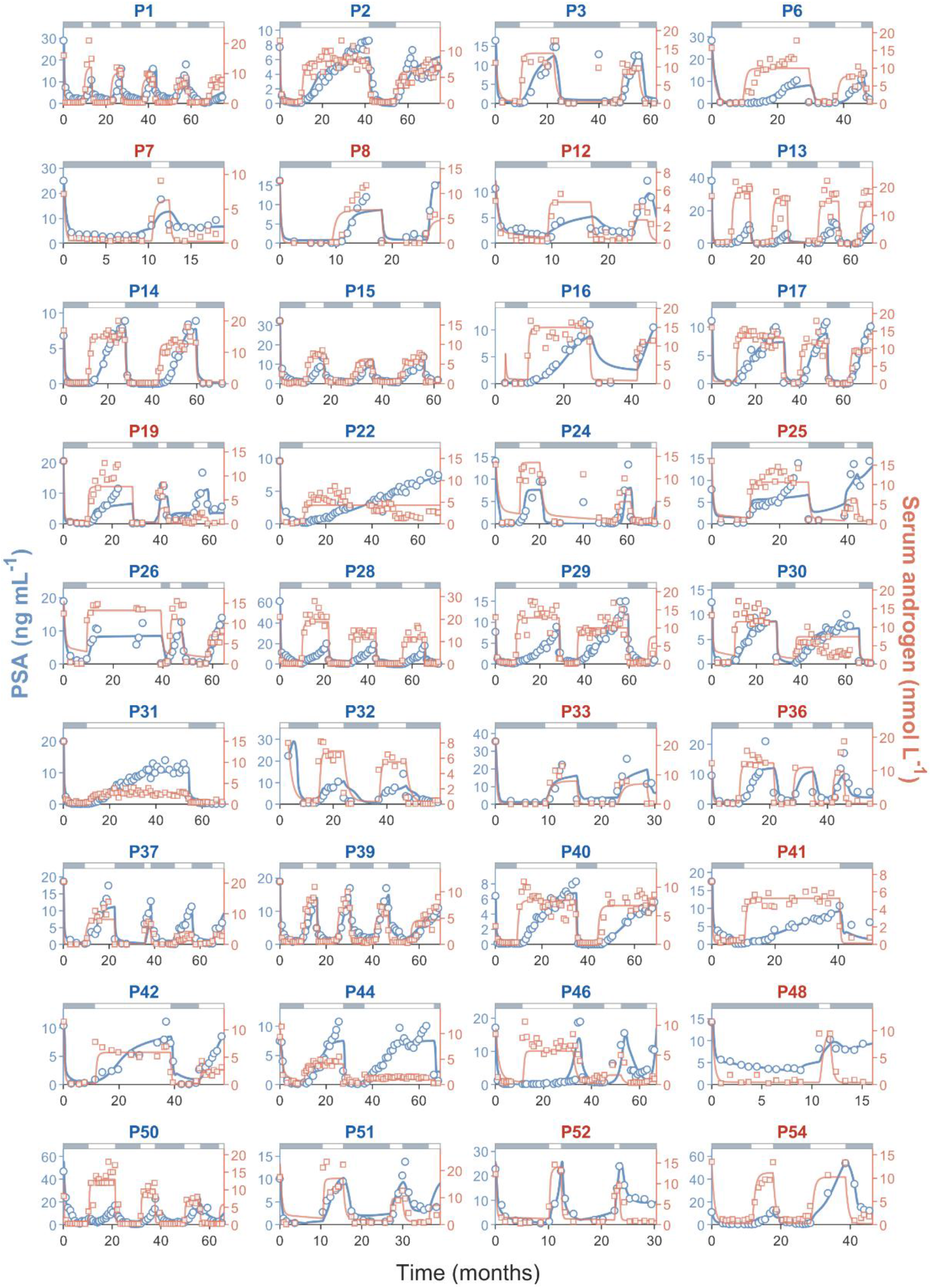
Model fit to Bruchovsky et al. data (continue in Fig. S2). Blue points/curves are PSA observations/model fit; Red points/curves are serum androgen observations/model fits. Grey bar on top indicates on treatment. White bar is off treatment. Blue subtitle indicates success. Red subtitle indicates failure. Companion to main-text Fig. 3A. Together, Fig. S1 and Fig. S2 cover all 71 patients with at least one complete cycle

**Fig. S2.**
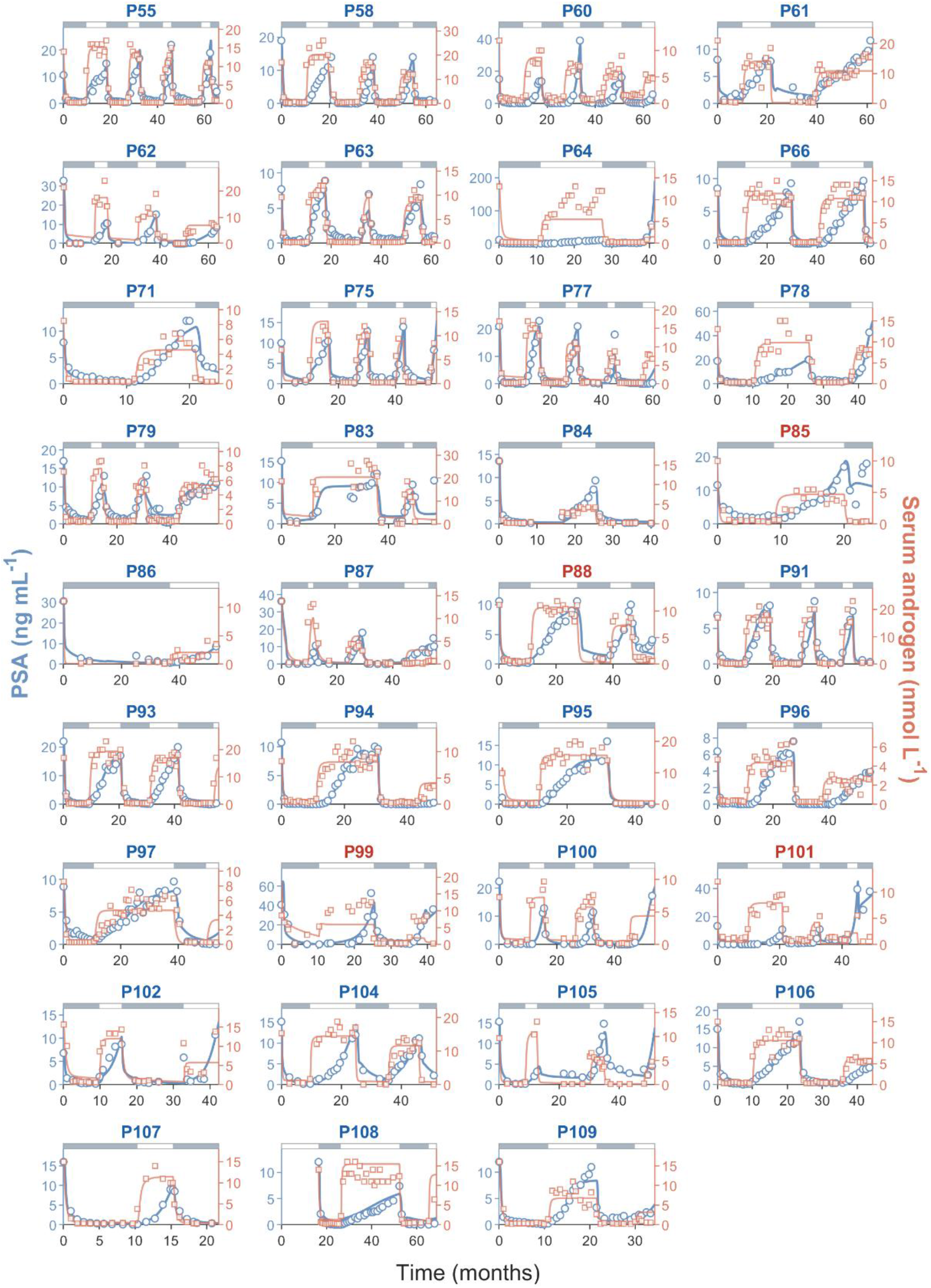
Model fit to Bruchovsky et al. data (continuation of Fig. S1). Blue points/curves are PSA observations/model fit; Red points/curves are serum androgen observations/model fits. Grey bar on top indicates on treatment. White bar is off treatment. Blue subtitle indicates success. Red subtitle indicates failure. Companion to main-text Fig. 3A. Together, Fig. S1 and Fig. S2 cover all 71 patients with at least one complete cycle.

**Fig. S3.**
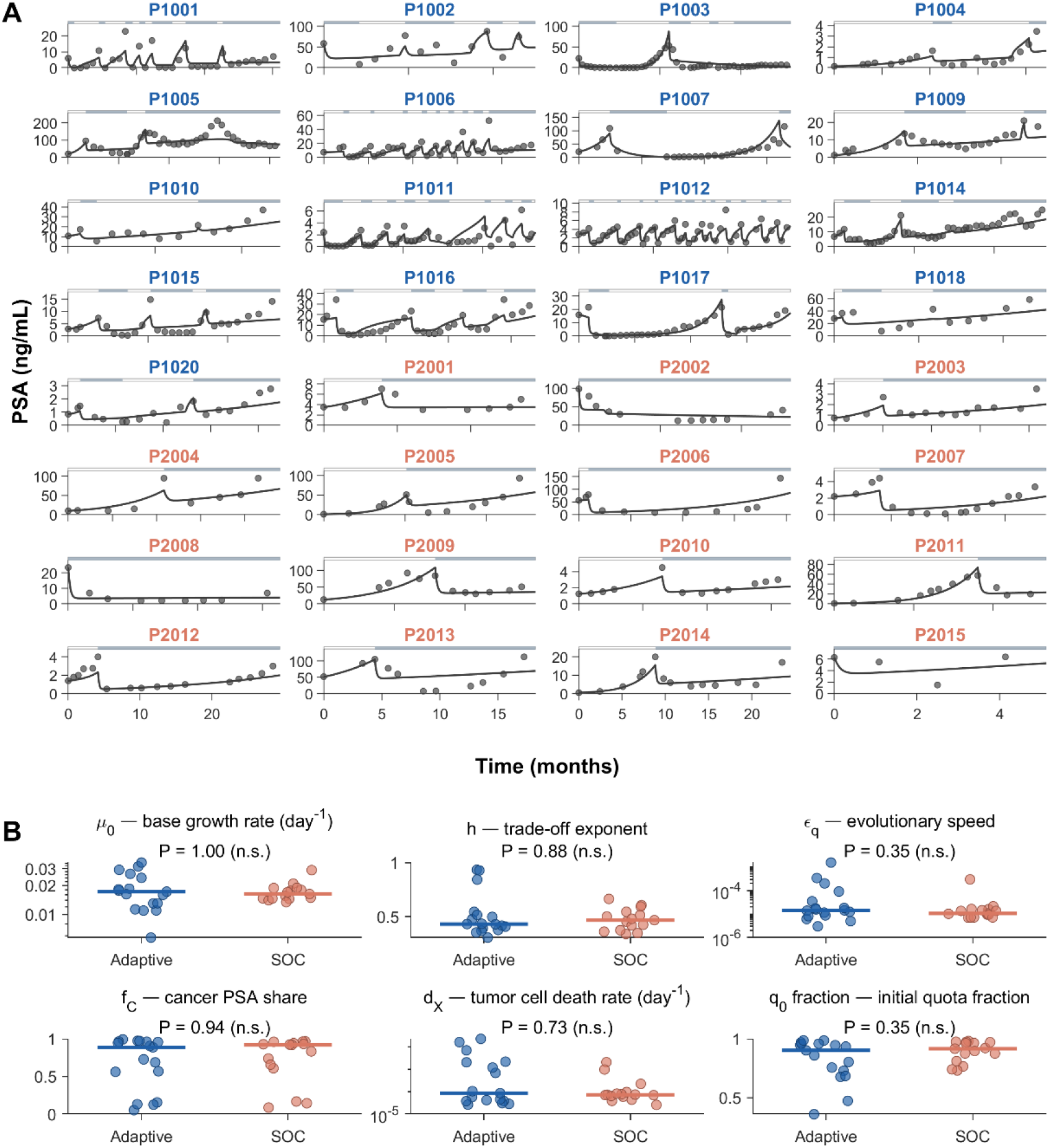
Simplified model fit to Zhang et al. data. (**A**) Blue points/curves are PSA observations/model fit. Grey bar on top indicates on treatment. White bar is off treatment. (**B**) Stratification of individual patients’ parameters based on the reported outcome. Mann-Whitney U-test are used for statistical comparisons. The blue vs. red subtitle indicate patients in the adaptive vs. standard-of-care arms, respectively.

**Fig. S4.**
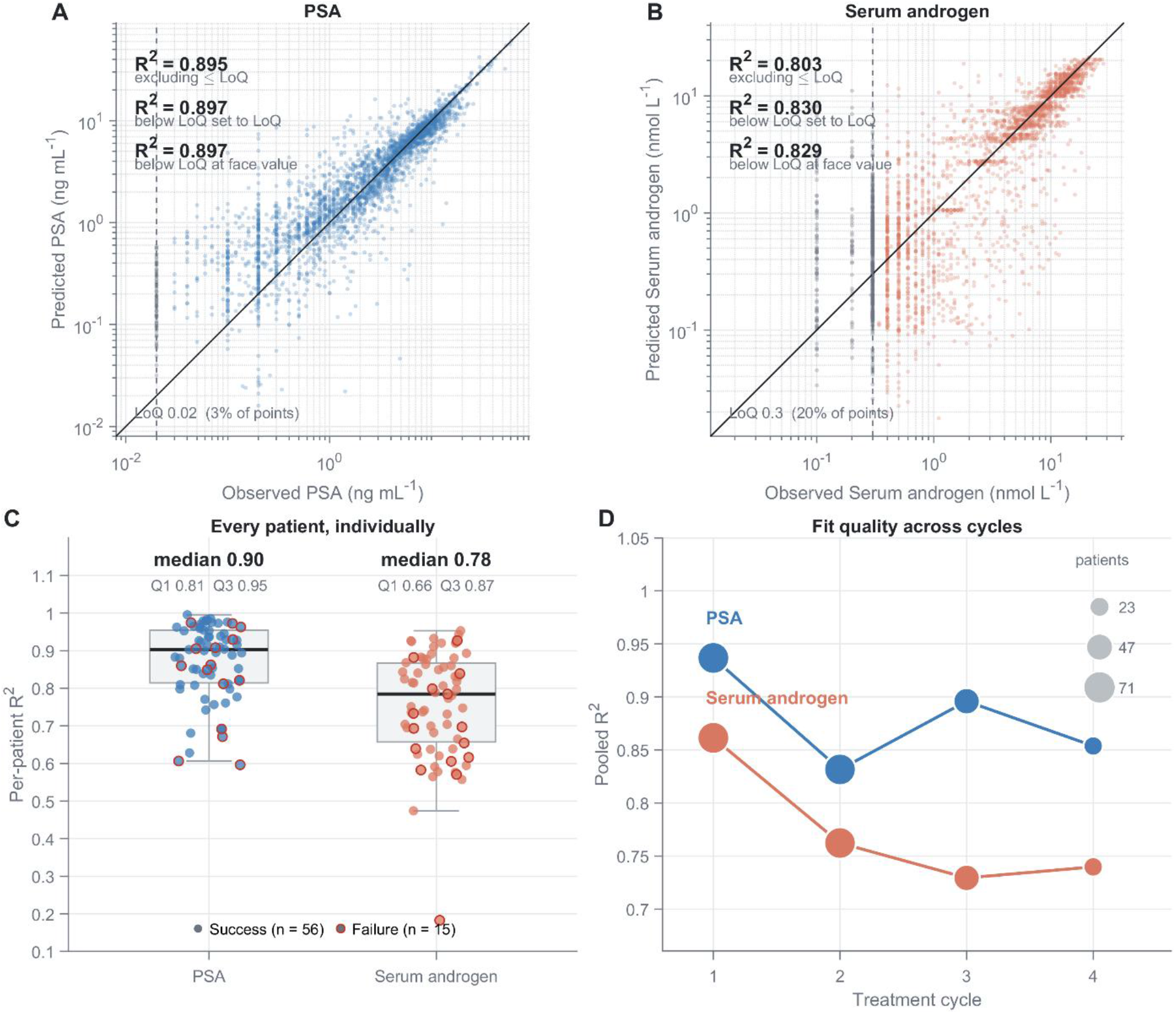
Agreement between model predictions and observed PSA and androgen levels for the Bruchovsky cohort. (**A–B**) Predicted (model best fit) versus observed PSA (3,057 measurements; 71 patients) and serum androgen (3,057 measurements). R^2^ was calculated excluding, censoring at, or retaining values below the assay floor (PSA: 0.02 ng/mL, n = 84; androgen: 0.3 nmol/L, n = 617; grey). (**C**) Patient-level R^2^ distributions. Treatment failures are outlined in red. Boxes show medians and interquartile ranges. Whiskers extend to 1.5 times interquartile-range whiskers. (**D**) R^2^ by treatment cycle (not cumulative). Dot size indicates the number of contributing patients. Fit quality was highest in cycle 1 and remained stable thereafter.

**Fig. S5.**
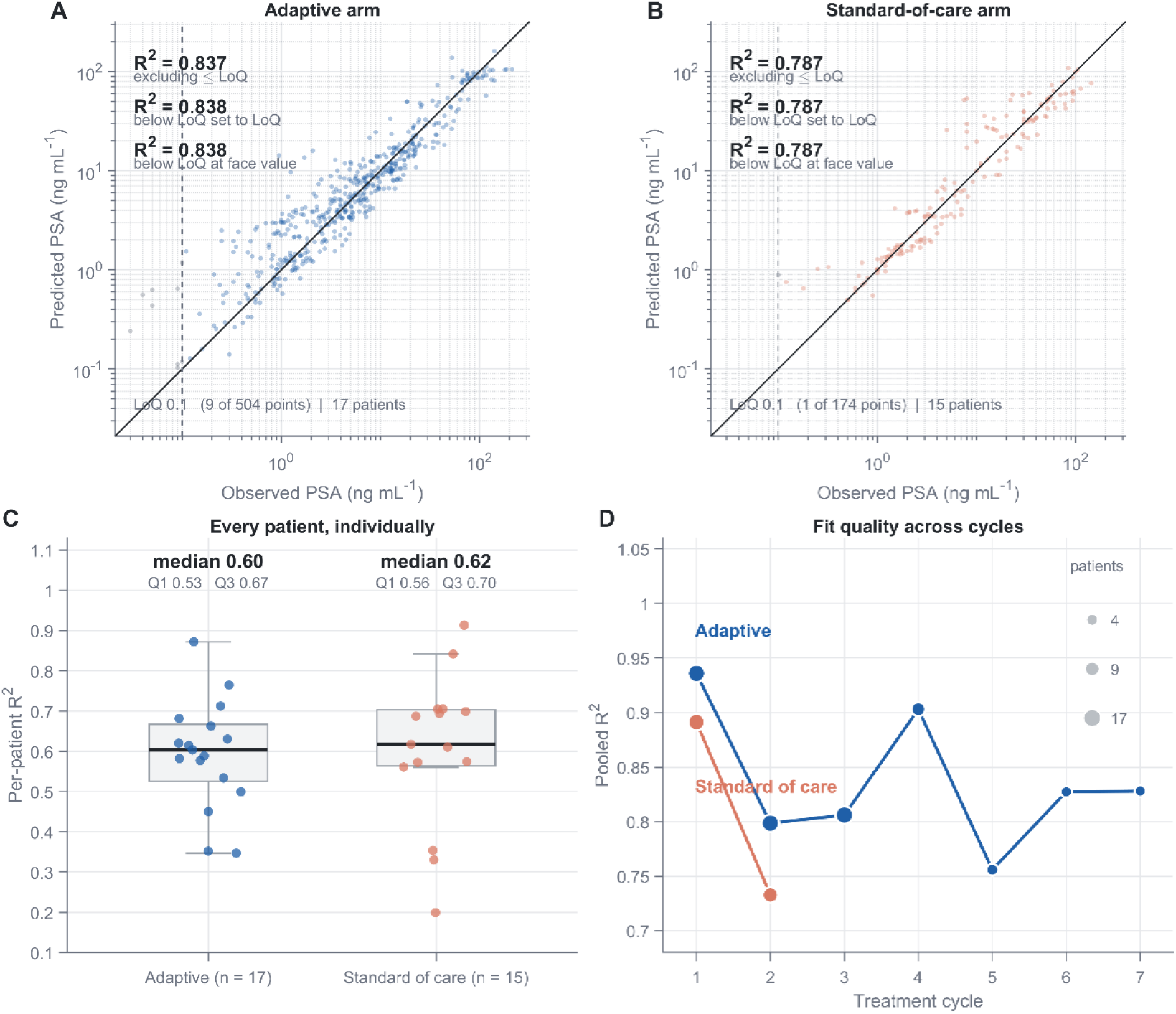
Agreement between reduced-model predictions and observed PSA levels in the Moffitt cohort. (**A–B**) Predicted (model best fit) versus observed PSA in the adaptive-therapy (17 patients) and standard-of-care (15 patients) arms. R^2^ was calculated excluding, censoring at, or retaining values below 0.1 ng/mL (n = 10). (**C**) Patient-level R^2^ distributions by treatment arm. Boxes show medians and interquartile ranges; whiskers extend to 1.5 times interquartile-range whiskers. (**D**) R^2^ by treatment cycle (not cumulative). Dot size indicates the number of contributing patients. Adaptive therapy is shown through cycle 7. For 12 out of 15 standard-of-care patients, cycles 1 and 2 represent the periods before and during continuous androgen deprivation, respectively. The other three (P2002, P2008, P2015) only has one cycle of continuous androgen deprivation.

